# Nitrogen acquisition and selective bacterial attraction through fungal hyphae promote plant growth and health under nitrogen limitation

**DOI:** 10.1101/2025.08.05.668771

**Authors:** Tan Anh Nhi Nguyen, Takuya Wada, Masami Nakamura, Yuniar Devi Utami, Salvatore Cosentino, Momoko Takagi, Kenta Ikazaki, Takatoshi Kiba, Kei Hiruma

## Abstract

Plants alleviate nitrogen (N) deficiency through interactions with beneficial microbes, yet mechanisms underlying plant-fungus-bacterium mediated plant growth promotion (PGP) remain poorly understood. Here, we show that the root endophytic fungus *Colletotrichum tofieldiae* (Ct) enhances the growth of diverse plant species beyond the Brassicaceae under N deficient field and lab conditions. Optimal Ct-mediated PGP requires *Arabidopsis thaliana* nitrate transporters (AtNRT2.1/2.2/2.4/2.5), as well as the fungal N utilization regulator CtAREA. Isotopic tracing with ^15^NO_3-_ reveals that Ct transfers N to hosts via CtAREA-dependent yet AtNRT2s-independent route, implicating distinct, parallel pathways in PGP. Moreover, Ct attracts the beneficial bacterium *Paraburkholderia monticola* through its hyphae, which metabolizes organic N sources and additively enhances plant growth with Ct. Notably, *P. monticola* suppressed the pathogenic transition of Ct in plant mutants defective in Brassicaceae-specific tryptophan-derived metabolism, potentially accounting for Ct’s broad beneficial activities. This plant-fungus-bacteria interaction may secure consistent PGP across varied agricultural settings.

---

Nitrogen (N) is an essential macronutrient for plant growth, but its availability is often limited in natural soils ^1^. While nitrate and ammonium are the main inorganic forms taken up by plants, organic forms such as amino acids in soils are generally inaccessible and may even inhibit growth ^2,3^. Given that nitrate is the preferred N source for many plant species ^4^, to efficiently acquire nitrate under N deficiency, plants have evolved intrinsic N starvation response (NSR) pathways, the activation of which enhances nitrate uptake through the induction of high-affinity nitrate transporters. In *A. thaliana*, while the dual-affinity transporter *NRT1.1* functions across a wide range of N concentrations, members of the high-affinity NRT2 family play a central role in NSR. *NRT2.1* and *NRT2.2* together account for the majority of nitrate direct uptake by plants under low-N conditions ^5^, whereas *NRT2.4* and *NRT2.5* are strongly induced under severe N starvation ^6,7^, collectively contributing to approximately 95% of high-affinity nitrate influx ^7^.

Despite these adaptive mechanisms, plants still face difficulties in acquiring N due to the limited range of rhizosphere. As an alternative strategy, plants form symbiosis with microbial partners, including fungi and bacteria, which assist in N acquisition and enable the utilization of otherwise inaccessible N sources. Among these microbial partners, fungi are particularly notable for their ability to extend hyphal networks through soil, effectively acting as extensions of plant roots. Beneficial fungi such as arbuscular mycorrhizal (AM) and ectomycorrhizal (ECM) fungi transfer N to the host ^8–12^. While N can be transferred in various forms, including nitrate and ammonium, most studies so far have focused on ammonium as the primary form of fungal-mediated N transfer ^11–14^. In contrast, few studies showed that nitrate was also reported to be up taken by AM fungi (AMF) and then transferred to the host ^12,15^. A recent study further showed that rice nitrate transporter *OsNPF4.5* is essential for AMF-mediated nitrate uptake, highlighting the potential roles of plant nitrate transporters in mediating beneficial plant–fungus interactions ^15^.

Unlike most land plants, many non-mycorrhizal plants, including members of the *Brassicaceae* family such as *Arabidopsis thaliana*, do not form AMF symbioses due to the lack the key gene sets ^16,17^. These plants instead form associations with other beneficial non-mycorrhizal fungi that help nutrient acquisition ^18–20^. However, direct evidence demonstrating that these fungi’s ability to transfer nitrate or related N to plants for plant growth under nitrate dominating conditions remains scarce. The knowledge gaps highlight the need to investigate how plants and their fungal partners coordinate nitrate uptake and metabolism during symbiotic interactions, beyond the classical AM symbioses.

In contrast to plants, which prefer nitrate as a N source, fungi preferentially utilize ammonium and glutamic acid ^21^. When these preferred sources are depleted, fungi activate secondary N utilization pathways to assimilate nitrate and nitrite. For example, in *Aspergillus nidulans* and *Neurospora crassa*, this process is globally regulated by the transcription factor Nit2/AREA, which, in conjunction with pathway-specific regulators such as Nit4 (for nitrate assimilation), controls the expression of downstream genes including nitrate reductase and nitrite reductase ^21–23^. Although the genetic basis for nitrate acquisition and utilization is well characterized in some model fungi, its role in beneficial plant–fungal symbiosis remains unexplored.

Despite the potential of beneficial fungi, most studies have been confined to laboratory or greenhouse settings, where environmental complexity is minimal. In the field, their effects are often inconsistent, possibly due to interactions with highly complex soil microbiota. On the contrary, recent studies imply the benefit of fungal hyphospheric bacteria. Similar to the rhizosphere, the surrounding of fungal hyphae contains hyphal exudates that influence the composition of nearby bacterial communities, often enriching taxa such as *Pseudomonas*, *Streptomyces*, and *Burkholderia* ^24^, which may enhance fungal symbiosis and potentially plant growth ^24–27^. For instance, addition of a soil with a microbial community have been shown to enhance AM fungi-mediated N mineralization from organic matter under N-limited conditions ^27^. These findings imply a synergistic role of fungi and associated bacteria in PGP. However, beyond the conceptual framework and preliminary observations, direct evidence demonstrating that hyphae-associated bacteria enriched in the rhizosphere contribute to plant growth in an additive manner and the underlying mechanisms, remain elusive.

*Colletotrichum tofieldiae* (Ct) is a non-mycorrhizal endophytic fungus that has been shown to colonize *A. thaliana* roots and transfer phosphorus to the host via its hyphal network under phosphate deficiency ^19^. Notably, Gonzalez et al., for example, reported significant PGP by the Ct strain 0861 in maize under fertilized-field conditions, although the underlying mechanisms remain unresolved. The result, along with similar studies using other endophytes, implies the potential of easily culturable, non-AM fungi as inoculants for promoting plant growth ^28^. Still, comprehensive studies that integrate both field and laboratory experiments simulating field conditions, alongside mechanistic investigation, remain markedly limited across all interaction models to date.

Here, we show that the root endophytic fungus *Colletotrichum tofieldiae* (Ct) promotes plant growth under N-limiting field and field-simulating laboratory conditions, in part by transferring N to the host and enhancing plant nitrate direct uptake by inducing plant nitrate transporters. The disruption of the fungal nitrate utilization pathway impairs Ct-mediated N transfer and PGP, highlighting its critical roles in the beneficial interaction. Furthermore, Ct presence alters the root-associated bacterial community, selectively enriching a bacterium that additively act with Ct to further promote plant growth, particularly when organic N sources are available. These findings provide the first genetic evidence of a fungal N utilization pathway indispensable for plant–fungus mutualism and identify a previously overlooked role of hyphae-associated beneficial bacteria in enhancing plant performance through a tripartite plant–fungus–bacterium interaction.

## Ct4 promotes Brassica plant growth in open fields and non-sterilized soil under N-deficiency

To investigate whether *C. tofieldiae* exhibit plant growth promotion (PGP) under field conditions, we performed a total of 9 open field trials in Japan. We utilized the native strain Ct MAFF 712334 (hereafter referred to as Ct4) ^29^ inoculated it onto *Brassica rapa* var. perviridis (Komatsuna in Japanese), a widely cultivated spinach in Asian countries. These trials were carried out in two geographically and climatically distinct regions of Japan: Nara Prefecture and Nagano Prefecture (Fig. 1a). Compared to Nara, Nagano, being the highest Prefecture in Japan, experienced a wider temperature ranges and lower annual rainfall (Fig. S1a). Both fields had previously been subjected to conventional agricultural practices involving chemical fertilization; however, no chemical fertilizers have been applied for at least the past five years. In Nara, field trials included both non-fertilized and fertilized plots. The former plots received no fertilizer for at least five consecutive years prior to the trial, whereas the latter plots were fertilized with a standard NPK fertilizer. In fertilized plots, no significant differences were observed between Ct4-treated and mock-treated plants (Fig. 1b). However, in non-fertilized plots, Ct4-treated plants exhibited significantly enhanced shoot growth compared to mock-treated plants (Fig. 1b). Soil analysis revealed that the non-fertilized field contained adequate phosphate (PO_43-_) for plant growth but was markedly depleted in soluble N, namely NO_3-_ and NH_4+_ (N ≤ 8 mg/100 g soil, Fig. S1b). These conditions were consistent across non-fertilized field trials, except at the Nara field in spring (Exp 3), where the N content were sufficient and no significant PGP was observed (Fig. S1c). Similarly, our Nagano field trials resulted in consistent Ct4-mediated PGP in non-fertilized fields, where N level was detected at a low level (Fig. 1c, Fig. S1c-d). In addition to *B. rapa*, Ct4 also enhanced the growth of *Lactuca sativa* (lettuce), a species distantly related to brassicas (Fig. S1e-f). These results suggest that despite climatic variations across seasons and geographic regions, Ct4 consistently promotes *B. rapa* shoot growth under N-limited conditions (Fig. 1d). Moreover, the observed PGP effects in lettuce suggest that the Ct4-plant beneficial interactions may be conserved across diverse plant lineages.

**Figure 1:**
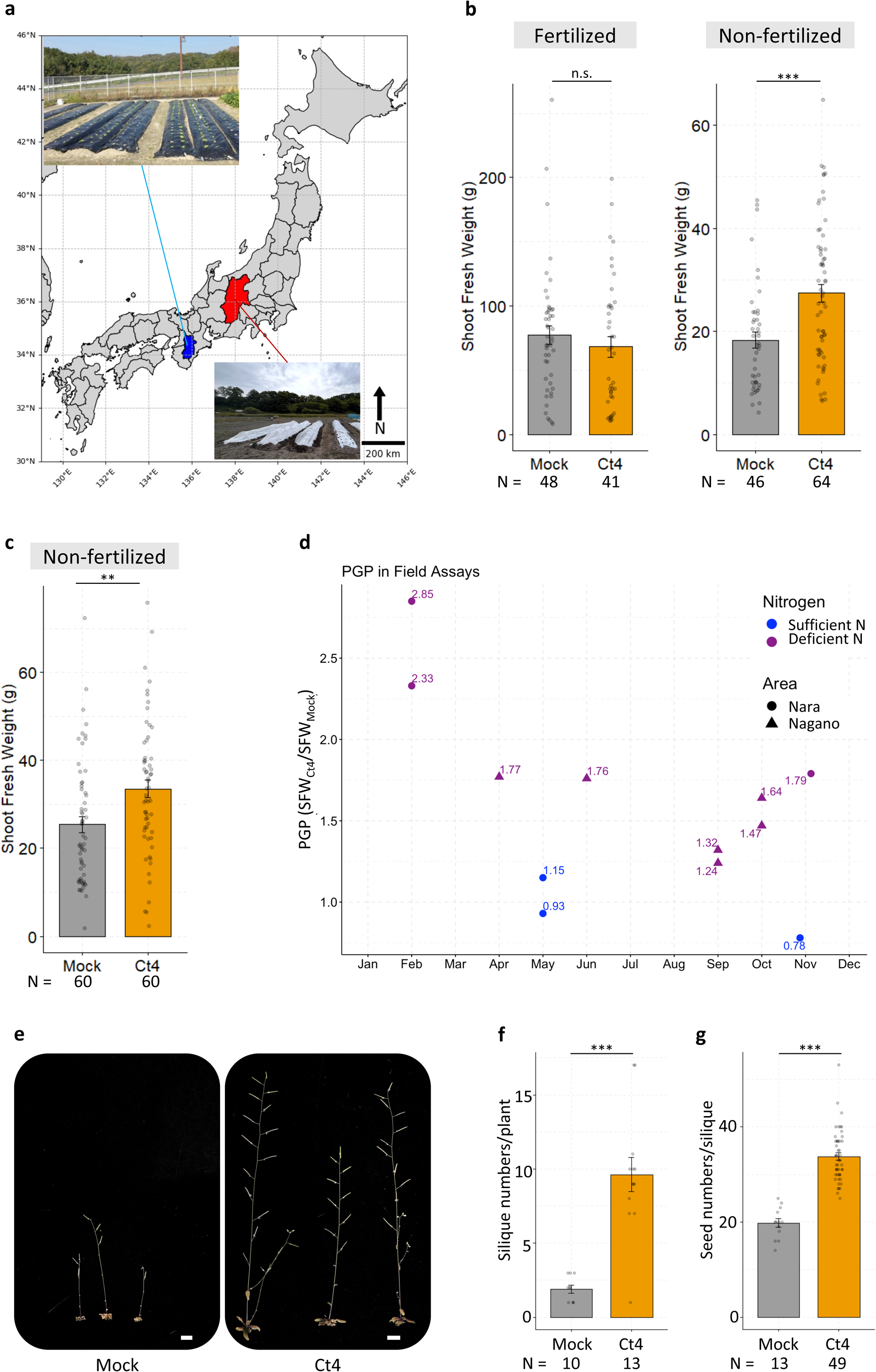
Ct4 promotes plant growth and fitness under low N conditions in open fields and non-sterilized soil pots. (a) Field assays were conducted in Nara (blue) and Nagano (red) prefecture, Japan. (b) Ct4-mediated PGP effects on *B. rapa* in Nara fertilized and non-fertilized fields (Exp 1, Fig. S1c), 30 day-post-inoculation (dpi). Mock, fertilized: N = 48; Ct4, fertilized: N = 41; Mock, non-fertilized: N = 46; Ct4, non-fertilized: N = 64. (c) Ct4-mediated PGP effects on *B. rapa* in Nagano (Exp 5, Fig. S1c), 29 dpi. N = 60. (d) PGP effects on *B. rapa* in field assays throughout different seasons in different years. Significant PGP can be observed when the nitrogen level in the soil was low (purple) but not when nitrogen was high (blue). (e-g) Ct4 promotes plant growth and fitness of *A. thaliana* plants in low nitrogen soil-pot assay. Bars = 1 cm. (f) Mock: N = 10; Ct4: N = 13. (g) Mock: N = 13; Ct4: N = 49. ** p < 0.01, *** p < 0.001, n.s. not significant, two-tailed t-test.

To further validate these field observations, we examined Ct4-mediated PGP under semi-controlled laboratory conditions, where Ct4 hyphae were mixed into non-sterilized soils, and *Arabidopsis thaliana* seedlings were cultivated in either Ct4-treated or mock-treated soils. Essential nutrients for plant growth, including phosphate, were supplied, whereas N sources (NO_3-_ and NH_4+_) were provided at minimal concentrations, mimicking nutrient limitation conditions observed in Nara and Nagano fields (Table S1). The *A. thaliana* plants grown in low N exhibited clear signs of stress (stunted growth, pale leaves), and the inoculation of Ct4 to the soil significantly promoted plant shoot growth (Fig. 1e, Fig. S2a). Such shoot growth enhancement was also observed in other plant species that we tested, including common crops such as Brassica plants (broccoli *B. oleracea*) and lettuce (*Lactuca sativa*) (Fig S2b-e). Notably, besides shoot growth enhancement, Ct4-treated plants displayed substantial increases in reproductive output, including a greater number of siliques and seeds of *A. thaliana* during the later stages of development (Fig. 1e-g). These field and laboratory assays demonstrate that Ct4 application markedly enhances plant growth and fitness under nitrogen-limiting conditions, extending beyond Brassica species.

## Ct4 colonization reprograms the transcriptome toward a symbiotic state, inducing the expression of plant nitrate transporters

We validated Ct4-mediated PGP observed in the field via inoculation assay using *A. thaliana* in a gnotobiotic agar system. This system allows precise control of each nutrient in both N-sufficient and N-deficient conditions, where nitrate-derived N comprised dominantly (See Methods and Table S1), and without any other microbial constituents (Fig. 2a). Ct4 consistently enhanced shoot biomass and chlorophyll content under N-deficient conditions, but not under N-sufficiency, a response correlated with increased root colonization observed specifically under N-deficient conditions (Fig. 2a, Fig. S3a-c). This suggests that Ct4-mediated PGP is dependent on N status. To investigate how plants respond to Ct4, we performed transcriptome analysis at early and established colonization stages under both N-sufficient and N-deficient conditions (Table S2). Microscopy analysis showed that Ct4 penetrated roots approximately 3 days after contact, with intensive colonization by 10 days (Fig. S4). As fungal contact occurred 3-4 days after spore placement in our inoculation assay, sampling at 6 and 15 dpi (∼3 and 12 days post-contact, respectively) captured transcriptomic changes occurring both prior to and following the onset of PGP (Fig. S5a–b). Principal coordinate analysis (PCoA) revealed distinct plant transcriptomic responses to N-deficiency and Ct4 colonization between 6 dpi and 15 dpi. At 6 dpi, mock-treated samples under N-sufficient and N-deficient conditions exhibited largely similar transcriptomic profiles. This similarity was also observed for Ct4 inoculated samples, although the transcriptomic profiles look different from those obtained for mock-treated samples (Fig. 2b). Notably, NSR marker genes *NRT2.1*, *NRT2.2*, *NRT2.4*, and *NRT2.5* were already induced under N deficiency at this stage (Fig. S5g, Table S2a), coinciding with a mild reduction in shoot biomass (Fig. S5a). These results suggest that while plants sensed N limitation at this stage, its physiological consequences were not yet pronounced. The Ct4-induced transcriptomic shift thus likely reflects early reprogramming toward beneficial associations rather than a direct response to N starvation. In contrast, at 15 dpi, mock-treated plants exhibited pronounced transcriptomic divergence between N-sufficient and N-deficient conditions, whereas Ct4-colonized samples showed a convergence of these profiles (Fig. 2b, Table S2d,f). This suggests that N starvation had a stronger impact on plants at this stage, but Ct4 colonization may have overridden the host’s starvation-induced responses, reprogramming the transcriptome toward a symbiotic state and mitigating the effects of external N limitation.

**Figure 2:**
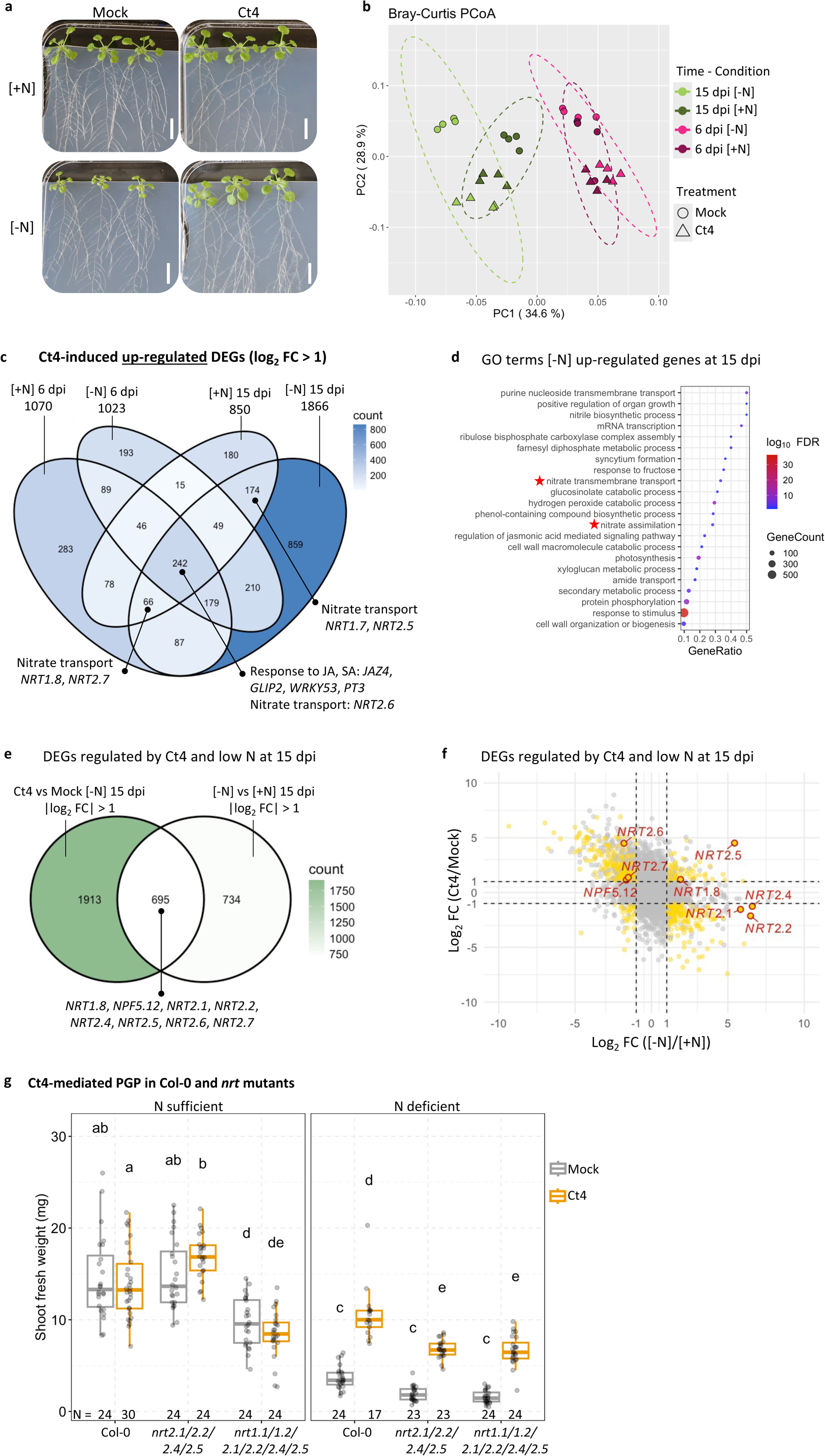
Ct4 induces plant’s nitrate transporters and promotes plant growth under low N conditions. (a) Ct4 promotes plant growth specifically under low N conditions. 7-day-old *A. thaliana* Col-0 plants grown in half-strength MS agar were transferred to [+N] or [-N] agar plates containing 11,400 µM and 275 µM N, respectively, with or without Ct4 inoculation. Bars = 1 cm. (b) Principal coordinate analysis (PCoA) of plant’s transcriptomic data from Col-0 root samples collected at 6 and 15 dpi. Colors indicate different time points and nutrient condition combinations (time-condition). Shapes represent treatments (circle: mock; triangle: Ct4). N = 4. Dashed-line circle: Distribution of each time-condition groups with 95% confidence. (c) Venn diagram showing the numbers of *A. thaliana* DEGs in response to Ct4 under N sufficiency [+N] and N deficiency [-N] treatment at 6 and 15 dpi. Numbers indicate DEGs counts. (d) GO terms enrichment analysis of 1,866 *A. thaliana* genes up-regulated by Ct4 under [-N] conditions (Fig. S5d) at 15 dpi (log_2_ FC > 1, FDR < 0.05). (e) Venn diagrams showing the numbers of *A. thaliana* DEGs in response to Ct4 or N deficiency at 15 dpi. Numbers indicate DEGs counts. (f) Scatter plot of DEGs regulated by Ct4 (y-axis) and by N deficiency (x-axis) at 15 dpi. Yellow dots indicate DEGs significantly regulated (|log_2_ FC| > 1, FDR < 0.05) by both Ct4 presence and N conditions. (g) Ct4-mediated PGP under N sufficiency [+N] and N deficiency [-N] agar plates (containing 11,400 µM and 275 µM N, respectively) in Col-0 and *nrt2.1/2.2/2.4/2.5* and *nrt1.1 nrt1.2 nrt2.1/2.2/2.4/2.5* mutants. Letters: post-hoc Tukey, p < 0.05.

Consistent with the PCoA, Venn diagram analysis revealed similar plant responses to Ct4 at 6 dpi across N conditions (Table S2b-c), with over 50% of Ct4-induced differentially expressed genes (DEGs) commonly upregulated under both N-sufficiency and N-deficiency (Fig. 2c, Fig. S5c, Table S2b-c). By contrast, at 15 dpi, Ct4 strongly impacted the plant’s transcriptome under N deficiency (1,866 DEGs), of which only ∼28% overlapped with those under N sufficiency (Fig. 2c, Fig. S5d, Table S2e-f). Despite this divergence, 242 DEGs were consistently upregulated by Ct4 across all conditions and were enriched in defense-related GO terms, including jasmonic acid and salicylic acid responses (Fig. 2c, Fig. S5e), suggesting the activation of a conserved plant response program associated with root colonization, independent of PGP. Given Ct4 up-regulated more plant’s genes under N deficiency at 15 dpi (1,866 DEGs) than the other treatment, where PGP was observed, we performed GO term enrichment analysis on these DEGs. The “nitrate transmembrane transport” GO term was significantly enriched, comprising nine NRT1s and NRT2s (Fig. 2d). These NRTs mediate both environmental nitrate uptake and source– sink nitrate redistribution ^6,7,30^, suggesting that Ct4 reprograms nitrate acquisition and allocation, which may mitigate plant responses to N starvation and promoting a transcriptional state resembling N sufficiency. We hence further investigated genes co-regulated by N starvation and Ct4 under N deficiency and found that nearly 50% of N starvation-responsive genes were also differentially regulated by Ct4 (Fig. 2e, Table S2d,f). Among these, eight *NRTs* were included, six of which displayed reversed expression patterns between N starvation and Ct4 inoculation: genes upregulated by N starvation were downregulated by Ct4 and vice versa (Fig. 2f). These include N starvation markers *NRT2.1*, *NRT2.2* and *NRT2.4*, whose expression was strongly induced by N starvation but shifted toward levels typical of N sufficiency upon Ct4 colonization (Fig. 2f, Fig. S5g-h).

To evaluate functional relevance of NRTs, we examined Ct4-mediated PGP in NRT mutants. *nrt2.1/2.2/2.4/2.5* mutants showed significantly reduced PGP under N deficiency but not under N sufficiency (Fig. 2g), aligning with their stronger transcriptional responses under low N. The sextuple mutants (*nrt1.1/1.2/2.1/2.2/2.4/2.5*) showed impaired growth under sufficient N but no further PGP reduction under N deficiency (Fig. 2g). Interestingly, even in the *nrt2.1/2.2/2.4/2.5* and *nrt1.1 nrt1.2 nrt2.1/2.2/2.4/2.5* mutants, Ct4-mediated PGP was not entirely abolished, implying that while these nitrate transporters facilitate PGP, they are not strictly essential, with *NRT1.1*, *NRT1.2* appearing to play only a minor role. Collectively, these results suggest that Ct4 colonization reprograms host transcription that contributes to plant nitrate acquisition against fluctuating N conditions, likely through modulation of NRT2-mediated pathways.

## *CtΔarea* -a Ct4 fungal mutant strain defective in N utilization-showed attenuated PGP

To identify fungal regulators involved in PGP under N deficiency, we conducted transcriptomic analyses of Ct4 and identified 16 up-regulated and 15 down-regulated DEGs in response to N starvation in *A. thaliana* roots (Fig. 3a, Table S3). Notably, expression of the fungal ammonium transporter *MEP3* and glutamine synthase *GS* was downregulated, while *NIT4* (nitrate assimilation transcription factor) and *NR* (nitrate reductase) were upregulated (Fig. 3a). This regulatory pattern aligns with the activation of the secondary N utilization pathway, which is regulated by the global regulator AREA and derepressed under ammonium/glutamine absence (Fig. 3b) ^21^. The metabolic shift from preferred to non-preferred N source utilization reflects fungal adaptation to low-N conditions and prompted us to examine the contribution of this regulatory pathway to Ct4-mediated PGP.

**Figure 3:**
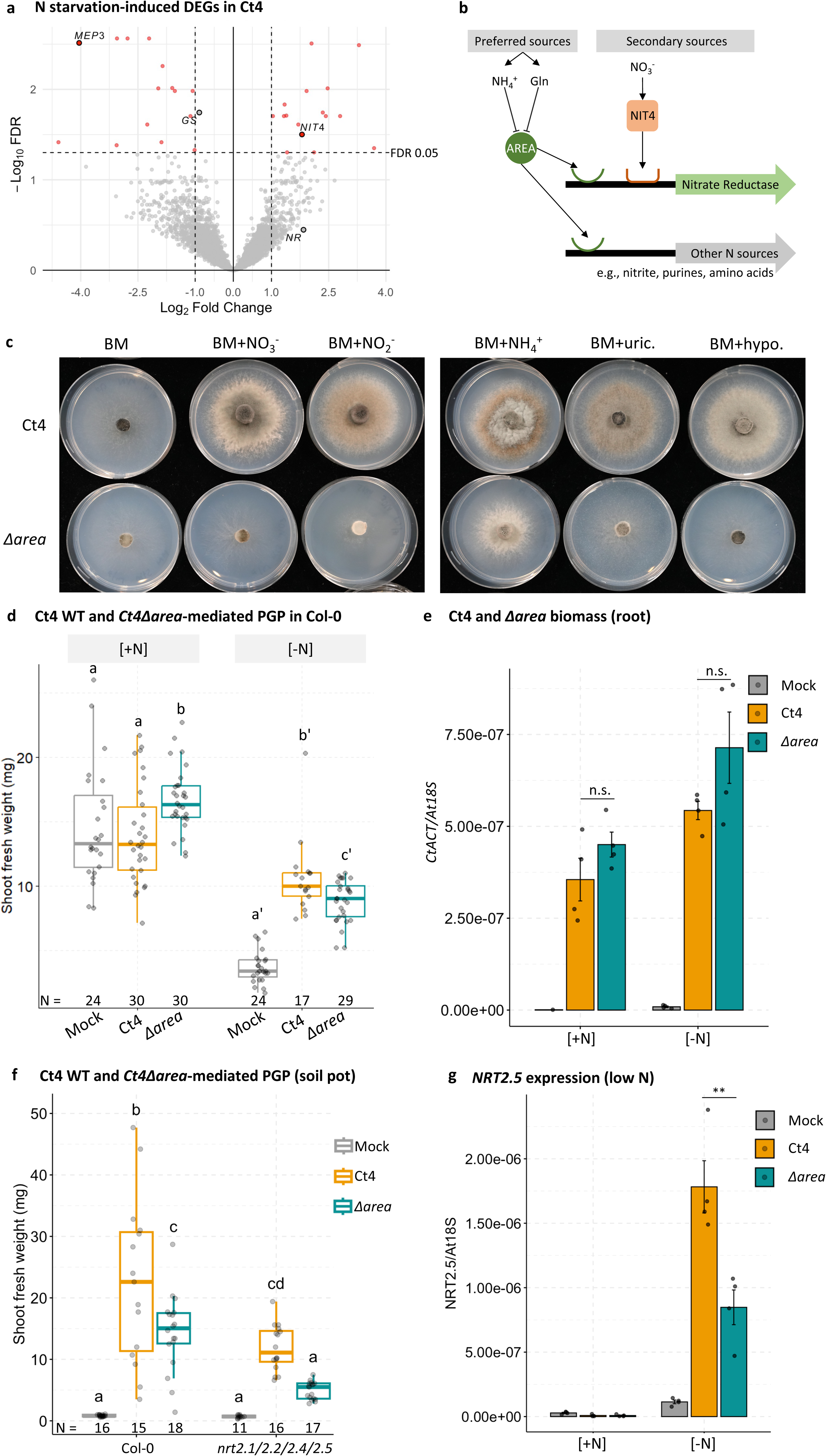
*CtAREA* regulates fungal nitrogen utilization and Ct4-mediated PGP under N deficiency. (a) Volcano plot showing Ct DEGs induced by N starvation in *A. thaliana* roots at 15 dpi. Non-significant genes are in grey; significant DEGs (|log_2_ FC| > 1, FDR < 0.05) are in red. (b) A schematic representation of the utilization of preferred (NH₄⁺ and glutamine) and secondary N sources in fungi. (c) Growth of wild-type Ct4 and *CtΔarea* mutants on a basal medium (BM), or a BM supplemented with the indicated N source. Equal-sized agar plugs either with Ct4 or *CtΔarea* were transferred to each medium and incubated for 5 days. Identical growth phenotypes were observed in four biological replicates. (d) Ct4- and *CtΔarea*-mediated PGP in Col-0 under N-sufficient and N-deficient conditions in agar system. (e) Ct4 biomass in the Ct4- and *CtΔarea*-inoculated roots, normalized to At18S in agar system. N = 4. (f) Ct4- and *CtΔarea* -mediated PGP in Col-0 and nitrate transporter mutants inoculation assay in soil pot assay. (g) *AtNRT2.5* expression level in the Ct4- and *CtΔarea* - inoculated roots in agar system, normalized to At18S. N = 4. Letters: post-hoc Tukey, p < 0.05. ** p < 0.01, *** p < 0.001, n.s. not significant, two-tailed t-test.

To this end, we generated a nitrate non-utilizing mutant (*nit2*) in the Ct4 background, via chlorate-induced mutagenesis method^31^. When grown on Mathur’s medium, a rich nutrient medium, *nit2* mutant’s growth or sporulation was comparable to WT Ct4 (Fig. S6a). However, on basal media (BM) supplemented only with nitrate, nitrite, or organic N (e.g., hypoxanthine) as N sources, the mutants exhibited reduced growth compared to Ct4 (Fig. 3c, Fig. S6b). This growth was largely rescued by adding ammonium, consistent with *nit2* mutants described in other fungi (Fig. 3b) ^32^.

The *nit2* defect has been reported to be attributed to mutations in the single-copy *NIT2*/*AREA* gene in other fungi^32–34^. Using NCBI BLAST, we identified a potential ortholog in Ct4, designated *CtAREA* (GenBank: GKT69611.1), which shares amino acid sequence similarity with *NIT2*/*AREA* from *Neurospora crassa*, *Aspergillus nidulans*, and *C. lindemuthianum* (Fig. S6c). Sequence analysis revealed a single-nucleotide mutation in the *CtAREA* gene region, where guanine nucleotide at position 2053 was deleted, resulting in a frameshift from amino acid 548 and a premature stop codon at position 697 (Fig. S6d). As AREA contains a Zn-binding domain spanning residues 659–709, this mutation led to the truncation of this domain in the *CtΔarea* mutants (Fig. S6d), which can cause the loss of function of AREA. Hence, we assigned *nit2* as *CtΔarea*. Inoculation assays in both agar and soil-based systems revealed that the *CtΔarea* mutants exhibited significantly reduced PGP effects under N-limiting conditions compared to Ct4 (Fig. 3d, Fig. S6e). Fungal biomass in roots colonized by Ct4 and *CtΔarea* was comparable (Fig. 3e, Fig. S6f), suggesting that the compromised PGP is not due to colonization failure. Notably, supplementing the soil pot with ammonium restored the growth-promoting effect of *CtΔarea* to WT levels (Fig. S6e), implying that *CtAREA* is required under conditions of severe N deficiency. Given the requirement of NRT2s for optimal PGP (Fig. 2g), we next investigated the role of CtAREA in NRT2-dependent PGP optimization. Inoculation of *nrt2.1/2.2/2.4/2.5* mutants with *CtΔarea* further reduced PGP, corresponded with a marked decrease in *NRT2.5* expression in Col-0 roots compared to those colonized by Ct4 (Fig. 3f–g). Conversely, constitutive expression of *NRT2.5* in *A. thaliana* Col-0 enhanced plant growth under N deficiency and partially rescued the *CtΔarea*-mediated defect (Fig. S6g). These results suggest that AREA-dependent fungal signaling contributes to *NRT2.5* induction in the host, which may contribute to PGP under N-limited conditions.

## Ct4 transfers N in a dependent manner of *CtAREA* but independently of root major *NRTs* and *AMT1;1*

We next aim to elucidate the roles of *AtNRT2* and *CtAREA* in Ct4-mediated PGP and to determine whether these components are functionally related. CtAREA governs nitrate utilization in fungi, and AtNRT2 transporters mediate nitrate uptake in plant roots. We therefore hypothesized that Ct4 may transfer nitrate to the host, with AtNRT2s facilitating its uptake, analogous to AM fungi, where a host nitrate transporter has been shown to regulate AM-mediated N uptake and growth promotion^15^. To address this, we employed a two-compartment assay, where ^15^NO₃⁻ were supplied exclusively in a compartment accessible only to fungal hyphae (HC), while ^14^NO₃⁻ were supplied in the shared plant–fungus compartment (PHC, Fig. 4a, Methods, Table S1). Additionally, we prepared a condition in which the PHC compartment was depleted of N (PHC = N0), thereby limiting AtNRT2s-mediating nitrate uptake pathway and making fungal-derived N from HC the predominant source supporting plant growth (Fig. 4a, Fig. 4b-c left panel). As a control, we included *Colletotrichum incanum* (Ci), a pathogenic fungus known to inhibit growth under phosphate limiting conditions and exhibit limited capacity to transfer phosphorus to host plants relative to Ct ^19^. Under N deficiency, however, Ci had no apparent effect on plant growth (Fig. S7a).

**Figure 4:**
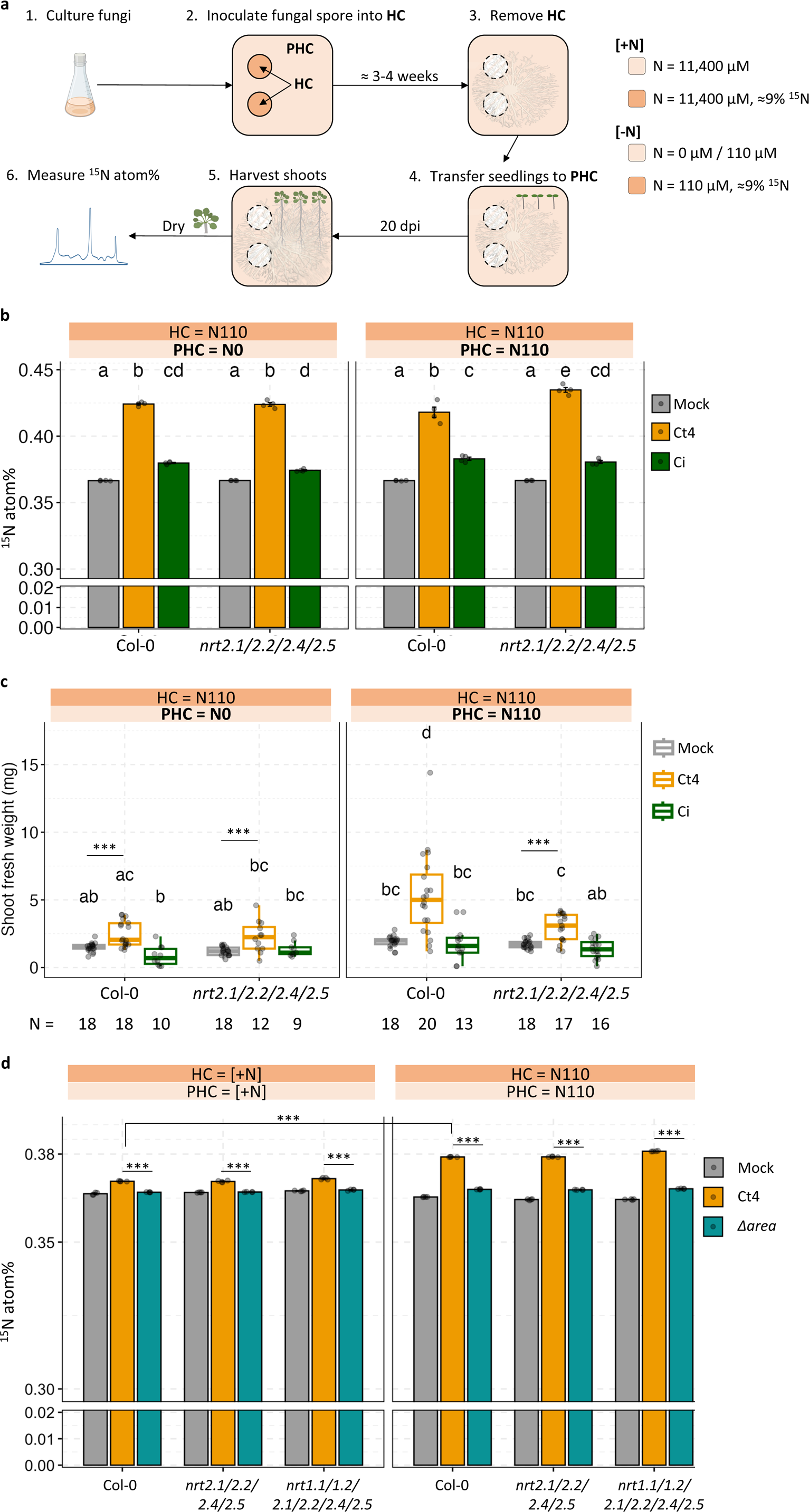
Ct4 transfers ^15^N to the host plants in a manner dependent on *CtAREA*, but independent of *AtNRT2s* and *AtAMT1;1*. (a) Experimental set up of ^15^N isotope 2-compartment assay. PHC: plant-hyphal compartment; HC: hyphal compartment. For N-sufficient conditions, both PHC and HC was supplemented with 11,400 µM of N (nitrate and ammonium); for N-deficient conditions, HC was supplemented with 110 µM of N, PHC was not supplemented with N (0 µM) or supplemented with 110 µM of N (110 µM). K^15^NO_3_ was used for HC, ^15^N atom% ≈ 9%. See also Table S1 for the details on the nutrient composition on PHC and HC. (b) ^15^N atom% in shoots of *A. thaliana* Col-0 and *nrt2.1/2.2/2.4/2.5* plants treated with DW (Mock), Ct4, or *C. incanum* (Ci) fungus under PHC = N0 and PHC = N110 conditions. N = 4. (c) Shoot fresh weight (mg) of *A. thaliana* Col-0 and *nrt2.1/2.2/2.4/2.5* plants at 20 dpi in PHC = N0 and PHC = N110 conditions in a 2-compartment assay. (d) ^15^N atom% in shoots of *A. thaliana* Col-0, *nrt2.1/2.2/2.4/2.5*, or *nrt1.1/1.2/2.1/2.2/2.4/2.5* plants treated with DW (mock), Ct4, or *CtΔarea* under N-sufficient or N-deficient conditions. N = 4. Letters: significant differences by post-hoc Tukey test, p < 0.05.*** p < 0.001, two-tailed t-test.

^15^N measurements revealed that, unlike mock- and Ci-treated plants, which exhibited ¹⁵N levels close to ¹⁵N natural abundance (∼0.365%), Ct4-treated plants displayed significantly elevated ¹⁵N enrichment (Fig. 4b), indicating direct N transfer from Ct4. The increase in ¹⁵N ratio was comparable between N0 and N110 conditions (Fig. 4b), implying that proportion of Ct4-mediated N acquisition is not substantially influenced by N acquired from the PHC compartment. Notably, under N0 conditions, both ¹⁵N atom% and PGP were similar between WT Col-0 and the *nrt2.1/2.2/2.4/2.5* mutants (Fig. 4b-c), indicating that Ct4-mediated N transfer from the HC compartment—and the resulting PGP—occurs independently of these NRT2 transporters. In contrast, under N110 conditions, despite comparable N transfer, PGP was significantly reduced in *nrt2.1/2.2/2.4/2.5* mutants (Fig. 4c), highlighting a critical role for these transporters in Ct4-mediated N acquisition from the PHC compartment. ¹⁵N ratio was also unaffected in *nrt1.1/1.2/2.1/2.2/2.4/2.*5 and *amt1;1* mutants, despite modest *NRT1.1*, *NRT1.2*, and *AMT1;1* induction by Ct4 (Fig. 4d, Fig. S5h, Fig. S7b-d). These results suggest that although Ct4 regulates the expression of several major and well-characterized host nitrate and ammonium transporters—some of which are critical for the plant’s intrinsic N uptake as well as Ct4-mediated PGP—the actual N transfer occurs independently of these transporters. Conversely, the ¹⁵N transfer was significantly diminished in plants colonized by *CtΔarea* mutants (Fig. 4d), resembling the phenotype observed in Ci-treated plants (Fig. 4b), despite Ci’s intact nitrate utilization capacity (Fig. S7e). This highlights the crucial role of *CtAREA* in Ct4-mediated ¹⁵NO^3-^utilization and subsequent N-transfer.

Under N-sufficient conditions, both NRT2-mediated PGP and Ct4-mediated N transfer were attenuated (Fig. 2g, Fig. 4c), likely reflecting a reduced plant demand for fungal-mediated N acquisition and explaining the minimal PGP observed under these conditions. Collectively, these findings suggest that Ct4 promotes host growth via CtAREA-dependent N transfer and induction of AtNRT2-mediated nitrate uptake, with these pathways functioning independently yet synergistically under N-limited conditions.

## Ct4 attracts plant growth promoting bacteria to roots under N-deficiency

Ct4 significantly promotes plant growth not only in axenic cultures but also in open-field and soil-pot assays, in the presence of root-associated microbial communities. Notably, under our tested conditions, the degree of Ct4-mediated PGP under non-sterile soil conditions often exceeds that observed in axenic agar systems, implying that Ct4 promotes plant growth through additional mechanisms beyond *CtAREA*-mediated direct N supply and activation of plant direct nitrate uptake. One possibility is that Ct4 modulates the root-associated microbial community in a way that promotes plant growth. Indeed, we observed a significant shift in the root-associated bacterial community of Ct4-inoculated plants under N-limiting field conditions (Fig. 5a). This shift was more pronounced in non-fertilized than in fertilized fields, indicating that Ct4 may have a stronger influence on shaping microbial communities under low-N conditions. In contrast, a distinct shift in the root-associated bacterial community was not observed in field assays where Ct4 failed to promote plant growth, specifically under conditions where soluble N was available in both fertilized and non-fertilized soils (Fig. S8a). To identify bacterial taxa enriched by Ct4 under low-N, we performed differential abundance analysis using ALDEx2 ^35,36^, which showed that Ct4 enriched more taxa in the root area under non-fertilized conditions (Fig. S8b, Table S4). We then investigated ASVs that showed significantly increased abundance in Ct4-inoculated roots in non-fertilized but not in fertilized fields (Fig. 5b, Table S5). Among these, we noted the enrichment of many ASVs belonging to *Burkholdericeae* and especially *Paraburkholderia* (Fig. 5b). Additionally, among the root-associated bacteria strains isolated from Ct4-treated plants, one *Paraburkholderia* strain, designated as Pam10, showed the strongest PGP effect under N-limited conditions containing amino acids, mimicking a soil condition (Fig. S10a-b) ^37^. Whole-genome sequencing revealed that Pam10’s 16S rRNA sequence matched ASV_88 (Fig. S9a), which was significantly enriched in Ct4-colonized roots only under non-fertilized, low-N conditions (Fig. 5b, Fig. S8c-d, Table S4-5). Pam10 taxonomy analysis using 16S rRNA-based and genome-based taxonomy revealed that Pam10 is a strain of *Paraburkholderia monticola* (Fig. 9b-d) ^38^

**Figure 5:**
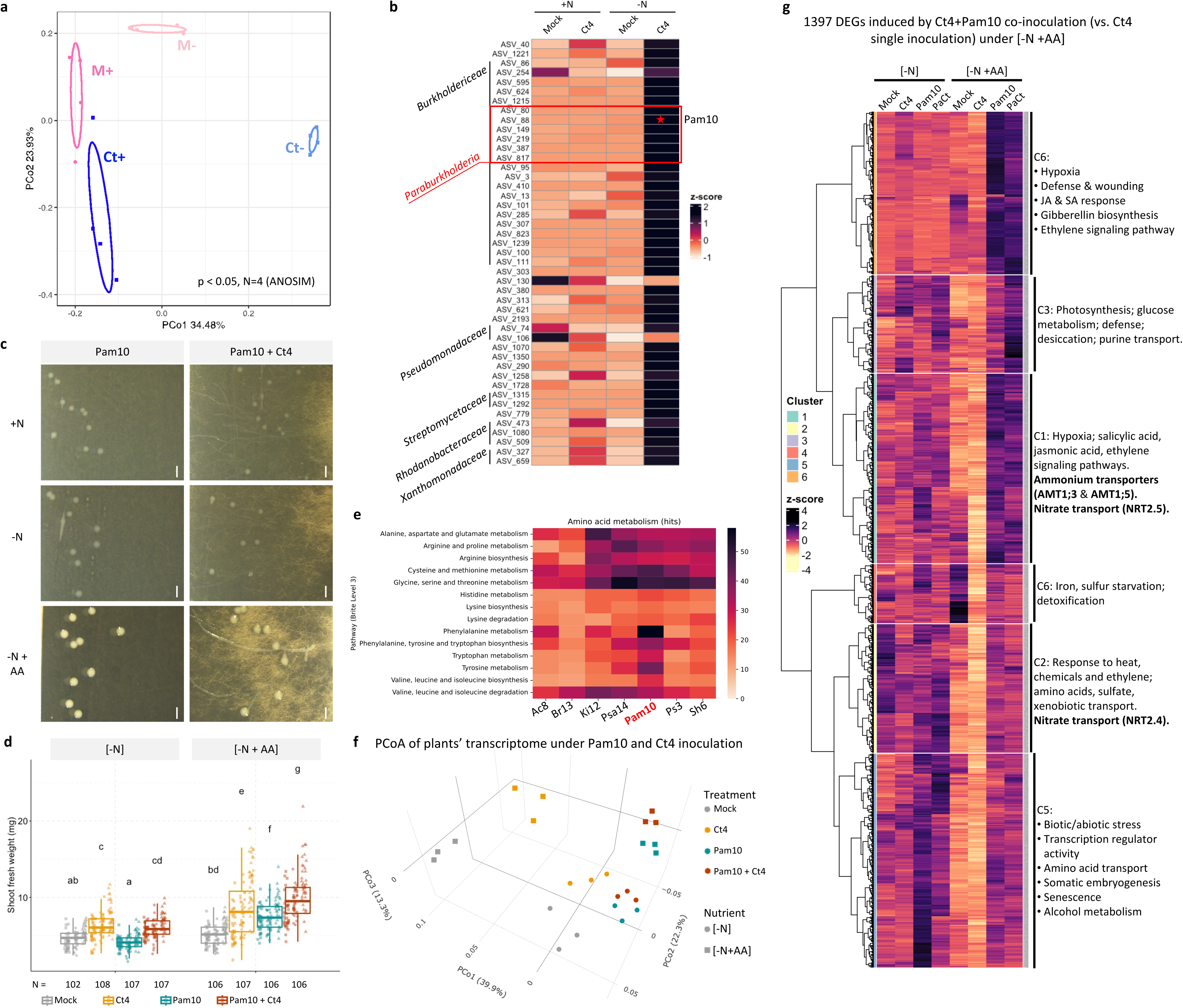
*Paraburkholderia monticola* (Pam10) isolated from Ct4-colonized roots additively promotes plant growth with Ct4 under specific N-limiting conditions. (a) PCoA of *B. rapa* root bacterial community profiles in fertilized (+N) and non-fertilized (−N) fields. M+, Ct+: Mock- and Ct4-treated root samples in fertilized fields; M−, Ct−: Mock- and Ct4-treated samples in non-fertilized fields. p < 0.05, ANOSIM. (b) Average relative abundance of bacterial ASVs enriched in *B. rapa* roots by Ct4 specifically in the non-fertilized, N-limited field, 4 samples/treatment. Enriched ASVs was chosen based on differential enrichment analysis by ALDEx2 ^35^. (c) Pam10 colonies with or without Ct4 under N-sufficient and N-deficient conditions with/without amino acids at 12 dpi by stereo microscope. Bars = 1 mm. (d) Inoculation assay on N-deficient agar with or without amino acids. Shoot fresh weight (mg) of *A. thaliana* inoculated with Mock, Ct4, Pam10 (OD₆₀₀ = 0.0001), or both, measured at 24 dpi. Pam10 and Ct4 co-inoculation showed additive PGP only with amino acid supplementation. Letters: significant differences by post-hoc Tukey test, p < 0.05. (e) Amino acid metabolism KEGG pathways analysis from genomes of bacterial strains isolated from *A. thaliana* roots colonized by Ct4 under N-limiting conditions. (f) PCoA from transcriptomic profiles (TPM) of plants in (d). 2D plots are in Fig. S11a-c. (g) Heatmap of z-scores (TPM) of 1,397 DEGs significantly upregulated in Pam10 + Ct4 co-inoculated roots vs. Ct4 alone under low-N + amino acid conditions (log_2_ FC > 1, FDR < 0.05). See also Table S8.

In culture, Pam10 formed compact, round colonies with limited spread when grown alone (Fig. 5c, Fig. S10c), suggesting their low mobility. This raised a question of how Pam10 becomes enriched in plant roots in the presence of Ct4. Some bacteria are known to utilize fungal hyphae as “highways” to access new niches, and recent studies have shown that AMF hyphae can bridge soil-dwelling PGPB to plant roots ^24,25^. We therefore examined whether Pam10 could utilize Ct4 hyphae to reach plant roots. Indeed, in the presence of Ct4, Pam10 spread along the fungal hyphae, forming a biofilm across both hyphae and root surfaces upon contact (Fig. 5c, Fig. S10d), suggesting that Ct4 recruits Pam10 to plant roots via its hyphal network. We then tested whether co-colonization with Ct4 and Pam10 enhances plant growth beyond that achieved by single inoculations. In a soil-pot assay, co-inoculation showed a clear additive PGP effect (Fig. S10e), whereas no such enhancement was observed under low-N conditions containing only inorganic N sources (nitrate and ammonium) in agar axenic culture (Fig. 5d). Importantly, genome analysis of Pam10 revealed that bacterial genes related to amino acid degradation were enriched compared with the other isolated bacterial strains (Fig. 5e), implying that Pam10 may utilize organic N sources. Correspondingly, Pam10 exhibited significantly enhanced growth when cultured with amino acid supplementation (valine, leucine, phenylalanine) (Fig. S10f-g), which exist in soils but were difficult for plants to efficiently utilize by themselves ^39^. Under these conditions, co-inoculation with Pam10 and Ct4 resulted in significantly greater PGP compared to either treatment alone, confirming an additive effect when organic N sources are present (Fig. 5d).

Transcriptomic analysis showed that Pam10 inoculation strongly altered plant gene expression under low-N conditions supplemented with amino acids (Table S6), with co-inoculation of Pam10 and Ct4 resulting in a transcriptomic profile closely resembling Pam10 single inoculation, indicating a dominant effect of Pam10 in this context (Fig. 5f, Fig. S11a-c). Supporting this, Venn analysis showed that Pam10 induced many *A. thaliana* genes under these conditions, with enrichment in the GO term “amino acid catabolic process” (Fig. S11d-e, Table S7). The co-inoculation of Pam10 and Ct4 triggered a distinct set of upregulated genes not observed in Ct4 single inoculation (Fig. S11f), enriched in nitrate, ammonium, and amino acid transport functions (Fig. 5g, Table S8). These patterns suggest that Pam10 may catabolize amino acids in the medium, supporting host uptake of inorganic N. Notably, this additive PGP effect was completely abolished in *A. thaliana nrt2.1/2.2/2.4/2.5* quadruple mutants, despite Ct4-mediated PGP in the mutants being comparable to that in wild-type plants under soil conditions (Fig. S10e), thereby, confirming that these NRT2s are required for the observed additive PGP. Taken together, these observations suggest that Ct4 attracts Pam10 to the rhizosphere under low-N conditions, enabling co-colonization at the plant root, which in turn facilitates the acquisition of both inorganic and complex N sources by host plants, thereby additively enhancing plant growth.

## Pam10 controls fungal growth under N-limiting conditions supplemented with amino acids

Interestingly, Pam10 also upregulated plant defense genes, particularly under amino acid-supplemented, N-deficiency conditions, both in the presence and absence of Ct4 (Fig. 5g, Table S8). These included genes implicated in systemic acquired resistance and antifungal responses, such as those involved in the tryptophan-derived metabolic pathway. It has been reported that Brassicaceae-specific tryptophan-derived metabolites synthesized by CYP79B2 and CYP79B3, such as camalexin and indole glucosinolates, play a pivotal role in limiting Ct4 overgrowth and preventing the transition of this beneficial fungus into a pathogenic state under phosphate-deficient conditions ^40^. Consistent with this, Ct4-mediated PGP was abolished in the *cyp79b2/b3* double mutant under N deficiency, with signs of disease on small leaves (Fig. 6a, Fig. S11g). Given that Pam10 robustly activated plant defense responses, particularly under N-deficient conditions supplemented with amino acids, we examined whether Pam10 could protect *cyp79b2/b3* double mutant plants from Ct colonization. Indeed, Ct4’s virulent symptoms on *cyp79b2/b3* mutants were suppressed when Pam10 co-existed, but only under conditions where Pam10 promoted plant growth ([-N + AA] condition, Fig. 6a, Fig. S11g). Consistently, Pam10 reduced Ct4 biomass significantly in both Col-0 and *cyp79b2/b3* roots under this condition (Fig. 6b). Moreover, when complex N sources were provided, Pam10 suppressed not only Ct4 but also the pathogenic fungus Ct3 (Fig. S11h-i), suggesting an antifungal activity independent of the host. Notably, despite successful suppression of Ct4 growth in *cyp79b2/b3*, additive PGP was not restored, suggesting that the tryptophan-derived metabolites are required not only to restrict Ct4 growth but also for the additive PGP.

**Figure 6:**
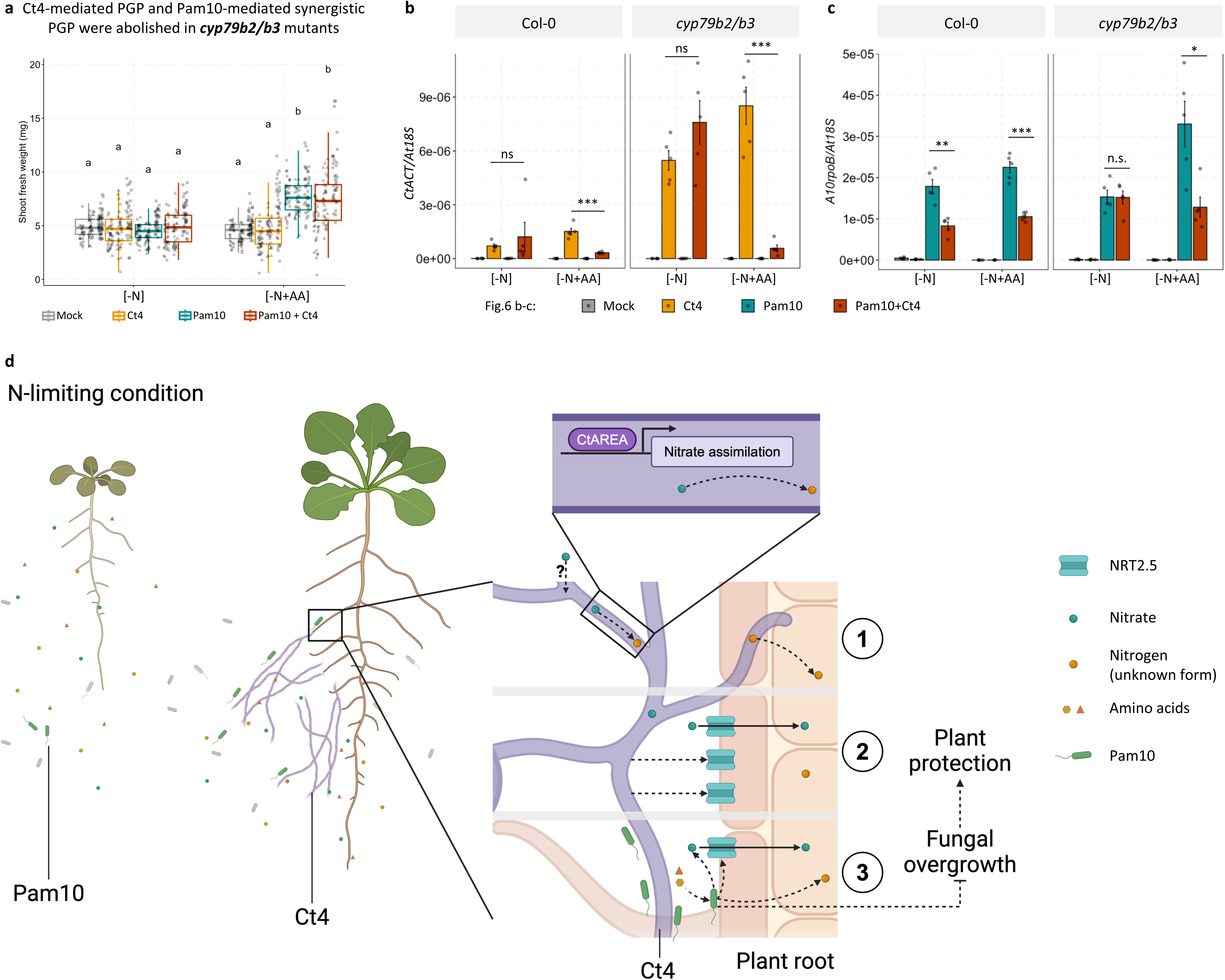
Pam10 protects immune-compromised *cyp79b2/b3* plants from Ct4 pathogenicity under amino acid-containing conditions and proposed model from this study. (a) Inoculation assay on *A. thaliana cyp79b2/b3* mutants under N-deficient conditions with or without amino acid supplementation. Shoot fresh weight (mg) measured at 24 dpi. Different letters indicate significant differences (one-way ANOVA, p < 0.05). (b-c) Quantification of fungal (b) and Pam10 (c) biomass in WT Col-0 and *cyp79b2/b3* mutants in single and co-inoculated conditions from Fig. 5e (Col-0) and Fig. 6a (*cyp79b2/b3*); normalized to plant 18S rRNA. N = 5. p < 0.05. * p < 0.05, ** p < 0.01, *** p < 0.001, n.s. not significant, two-tailed t-test. (d) Proposed model: Under low-N conditions containing both inorganic and complex N sources (e.g., amino acids), plant growth is limited by poor access to nitrate. Ct4 promotes PGP by: (1) transporting nitrate from otherwise inaccessible zones, via a nitrate utilization pathway regulated by the global regulator CtAREA; (2) facilitating nitrate uptake by enhancing plant *AtNRT2.5* expression, partially involving CtAREA; and (3) selectively attracting rhizobacteria (e.g., Pam10) that metabolize complex N sources and convert them into plant-available forms (e.g., nitrate). This coordinated colonization also helps restrain microbial overgrowth and reinforces plant defense responses.

Unexpectedly, unlike the case of the Ct4 hyphosphere (Fig.5c), Ct4 suppressed Pam10 proliferation on plant roots, suggesting reciprocal growth control between the two. This control was disrupted when plant immunity was compromised in the *cyp79b2/b3* mutant and amino acids were absent, a condition under which Pam10 failed to confer PGP and Ct4 showed virulence (Fig. 6c). The diminished PGP under this condition suggests that the balance maintained by plant immunity and Ct4–Pam10 reciprocal control is essential for PGP in co-inoculated plants. Together, these findings suggest that Pam10 maintains plant health by enhancing plant defense and restraining fungal growth, while Ct4 concurrently restricts Pam10 proliferation. Such reciprocal inhibition in plant roots may serve as a regulatory mechanism to stabilize the tripartite interaction and optimize plant growth under N-limited conditions.

## Discussion and Conclusion

In this study, we report that the endophytic fungus *Colletotrichum tofieldiae* strain Ct4 promotes plant growth in both *Brassicaceae* and non-*Brassicaceae* species under N-limiting conditions in both laboratory and open-field environments, where complex environmental factors such as organic N sources and microbial communities are present. We propose a model for the underlying mechanisms of this PGP in Fig. 6d, involving both direct and indirect pathways. First, Ct4 transports nitrogen from otherwise inaccessible zones for plants via a nitrate utilization pathway regulated by the global regulator *CtAREA* (Fig. 6d (1)). Second, Ct4 enhances plant nitrate uptake by inducing *AtNRTs* (e.g., *NRT2.5*), a process that also involves *CtAREA* (Fig. 6d (2)). Third, Ct4 selectively enriches beneficial rhizobacteria such as Pam10 in the rhizosphere (Fig. 6d (3)). The bacteria metabolize complex N sources and convert them into plant-assimilable forms (e.g., nitrate). Additionally, Pam10 contributes to microbial homeostasis by restricting potential fungal overgrowth, directly and possibly indirectly via activation of plant immunity. These complementary, and potentially synergistic, mechanisms can collectively underlie the robust PGP consistently observed in field and soil-based assays, where plants interact with diverse nutrient forms and microbial consortia.

We identify fungal secondary N utilization, regulated by the transcription factor CtAREA, as essential for fungal nitrate assimilation, N transfer to the host, and subsequent Ct4-mediated PGP under N-limiting conditions—representing the first documented fungal genetic basis for fungal-mediated PGP in such an environment. Notably, the *CtΔarea* mutants colonized plant roots at levels comparable to wild-type Ct4, suggesting that the diminished N transfer and PGP are not attributable to impaired root colonization. This aligns with findings in the pathogenic fungus *C. lindemuthianum*, where *ClAREA* disruption did not affect the early stages (i.e., penetration and initial biotrophic phases) of the infection but instead affected the later necrotrophic stages involving host cell death ^32^. Although the precise mechanism by which the *CtΔarea* mutants fail to transfer N remains to be elucidated, CtAREA-mediated pathway appears to be critical for nitrate assimilation after nitrate uptake, particularly under conditions where nitrate is the predominant nitrogen source—a deficiency that can be compensated by ammonium supplementation (Fig. S6e). Given the conservation of AREA and its regulatory functions across filamentous fungal taxa ^21,32,33^, it is potential that this regulatory pathway plays a similar role in other plant-associated fungi.

Importantly, the N transfer observed in Ct4 is absent in the closely related species *C. incanum* (Fig. 3g) which failed to promote plant growth under the tested N-limiting conditions despite possessing a functional N assimilation pathway (Fig. 4b-c, Fig.S7e). This suggests that the mechanisms underlying N transfer, and the resulting PGP may be attributed to relatively subtle genomic distinctions beyond *AREA* between Ct and *C. incanum* ^41^. A similar specificity has been observed in Ct-mediated phosphorus transfer^19^, also highlighting the potential use of *C. tofieldiae* as a model for studying nutrient exchange and signaling in plant-fungal beneficial interactions.

Recent studies identified the ammonium transporter ZmAMT3;1 and the nitrate transporter OsNPF4.5 as key components in AMF-mediated N uptake ^14,15^. In *A. thaliana*, the two transporters that share highest sequence similarity with *ZmAMT3;1* and *OsNPF4.5* are *AtAMT2* and *AtNRT1.2*, respectively. However, under our tested conditions, *AtAMT2* did not show a clear increase in transcript levels (Fig. S12a), and disruption of *AtNRT1.2*,along with its closely related *AtNRT1.1,* did not impair fungal N transfer (Fig. 3d), implying mechanistic differences between AMF and Ct symbioses. Although major root nitrate uptake transporters in *A. thaliana* were not essential for Ct4-mediated N transfer, they contribute to PGP under N-deficiency (Fig. 3e). Among them, *AtNRT2.5* was strongly upregulated by Ct4 regardless of N availability (Fig. S4h), and this induction was compromised in *CtΔarea* mutants. The overexpression of *AtNRT2.5* partially rescued the compromised PGP phenotype in *CtΔarea* in soil-grown plants (Fig.4f). These findings suggest that Ct4, via CtAREA, not only transfers N to the host but also enhances the host direct nitrate uptake. This, once again, highlights the difference with AMF-mediated nutrient acquisition, where nutrient uptake via mycorrhizal pathway antagonizes host’s direct uptake pathway ^42^. Given the conservation of *AREA* in filamentous fungi and *NRT2.5* in plants, further investigation into their coordinated roles under N-limiting conditions may shed light on how nitrate signaling and transport contribute to fungus-plant symbiosis.

Beyond the well-characterized plant–fungus bipartite symbiosis, increasing attention has been given to the role of bacteria in shaping plant-fungus-bacteria multipartite interactions. In particular, hyphospheric bacteria can mediate the host’s utilization of complex N sources ^43^. Our findings on plant-Ct4-Pam10 tripartite interaction support this emerging concept. Specifically, Pam10 benefits from Ct4 hyphal networks to reach the plant root, while in return enhancing host growth under N-limited conditions. Notably, Ct4 and Pam10 exhibit reciprocal control of their colonization in roots especially when they show PGP, suggesting a form of microbial growth restraint that maintains symbiotic balance and prevents over-colonization. The host’s defense via tryptophan-derived metabolism pathway also drives both parties’ growth, simultaneously restricting fungal growth and enabling Ct4 to suppress the Pam10 growth (Fig. 6b-c), a dynamic that unexpectedly results in maximum PGP under the tested conditions. This points to a regulatory mechanism where all partners actively coordinate each other’s growth to maintain a stable mutualistic interaction that supports plant growth under nutrient limiting conditions, although further investigation is required to address how this relationship benefits the associated microbial community.

Finally, although plant–beneficial fungal interactions have been extensively studied, translating these beneficial effects from controlled laboratory settings to field conditions remains challenging ^44^. Conversely, studies demonstrating significant PGP in field have often lacked mechanistic insight at the molecular level. In this study, we bridge this gap by linking the robust PGP conferred by a beneficial fungal endophyte under field conditions to a potential molecular mechanism elucidated under laboratory settings. Our recent findings demonstrate that constitutive expression of a fungal transcription factor, *CtBOT6*, converts this otherwise beneficial fungus into a pathogen under axenic conditions ^40^. This underscores the pivotal importance of uncovering the mechanisms by which beneficial fungi attenuate their intrinsic virulence, potentially through interactions with rhizosphere microbiota—as exemplified by the protective role of Pam10 against Ct4 over-proliferation in immuno-compromised *A. thaliana cyp79b2/b3* mutants. Elucidating these mechanisms represents a critical step toward the strategic deployment of such microbes in sustainable agricultural practices. To this end, our findings provide a model for investigating nutrient exchange within tripartite plant–fungus–bacterium interactions along the mutualistic–parasitic continuum, an intrinsic feature potentially shared by all microbial partners.

## Acknowledgements

We thank Akemi Uchiyama, Hiromi Haba, and Tokuko Sekine for their technical assistance. We thank Yasuyuki Kubo and Sayo Kodama for the published materials. We thank Yusuke Saijo and Shigeyuki Moteki for help to use fields in Nara and Nagano, respectively. We thank Klaus Schläppi, Yasuhiro Tsujimoto for fruitful discussion and help.

## Author contributions

KH initiated the project. NN and KH directed the research. NN, TW, MN, YU, MT, KI, TK and KH conducted the experiments. NN and KH conducted RNA-seq analyses. SC conducted comparative bacterial genome analyses. NN, TW, MN, YU, SC, MT, KI, TK and KH analyzed the data. NN and KH wrote the manuscript with feedback from all authors.

## Data and materials availability

Raw sequencing data of RNA-seq transcriptomic data have been deposited in DDBJ (PRJDB18752 for Fig.2 and Fig.3, PRJDB35609 for Fig.5). Raw sequencing data of 16S meta-sequencing data have been deposited in DDBJ (PRJDB35688). Bacterial genomes have been deposited in DDBJ (PRJDB35513). The Python code to perform the gene prediction, KEGG annotation, and to generate the heatmaps is available at https://gitlab.com/salvo981/genome_data_analysis.

## Materials and Methods

### Plant material and growth conditions

*Arabidopsis thaliana* plants used in this study were Col-0 background. The quadruple mutant *nrt2.1 nrt2.2 nrt2.4-1 nrt2.5* (referred to as *nrt2.1/2.2/2.4/2.5*) ^7^, and the sextuple mutant *chl1-12 nrt1.2 nrt2.1 nrt2.2 nrt2.4 nrt2.5* (referred to as *nrt1.1/1.2/2.1/2.2/2.4/2.5*), were generated via standard cross. The *amt1;1* mutant (CS6168), the *cyp79b2/b3* ^45^, and the *A. thaliana* expressing PIP2A-mCherry ^19^ were used in this study.

Seeds were surface sterilized with 70% ethanol for 30 seconds, followed by 6% sodium hypochlorite with 0.01% Triton X. After being washed five times in sterilized water, seeds were sowed on half-strength Murashige and Skoog (MS) vertical agar plates without sucrose addition (pH 5.8) at 22°C under fluorescent light (100 µmol m^−2^ s^−1^, 16 h light/8 h dark) for 7 days before inoculation.

### Fungal inoculation for pot and field assays

The fungal-culturing substrate was prepared by mixing 180 g sawdust, 60 g rice bran, 60 g wheat bran, and 180 mL water. The mix was divided (40-50 g substrate/box) into autoclavable plant box with ventilation hole (BMS, BC-PB851), plugged with cotton plug and sterilized by autoclaving twice at 121°C for 30 minutes. The substrate was cooled to room temperature before use. Fungal spores cultured in Mathur’s flask (described below) was harvested and resuspended in 10 mL distilled water (DW). This spores suspension or DW (mock treatment) was then thoroughly mixed into the substrate and incubated at room temperature for 7 days. The treated substrates-either with or without fungal mycelia-were subsequently combined with a soil matrix composed of 400 g vermiculite (LIFELEX, 0.15 kg/L), 100 g black soil (KOHNAN SHOJI Co., Ltd.), and 25 g of the prepared substrate, then divided into planting pots (diameter 7.5 cm, height 6.8 cm, Kyofuku). Two 7-day-old seedlings of *A. thaliana* were transplanted per pot, or two seeds of *B. rapa* (komatsuna, Sakata, Misaki 922770) or one seed of *Lactuca sativa* (lettuce, TOHOKU SEED Co., Ltd, noble SP 03514) were sowed directly. Plants were cultivated in a semi-controlled plant room (22°C, 16 h light/8 h dark, around 40% humidity), watered with tap water every 2–3 days.

For pot assay, plants were kept growing in the same conditions and irrigated weekly with approximately 30-40 mL (per pot) of either N-sufficient (11.4 mM) or N-deficient (110 µM or 275 µM) half-strength MS liquid solution (see below for the recipe) and additionally watered with tap water every 2–3 days until harvest.

For field assays, plants were transplanted into field plots after 7 days (*A. thaliana* and komatsuna), or 14 days (lettuce) for continued growth under natural field conditions, spaced at 15 cm intervals for *B. rapa* and 30 cm for lettuce. For field fertilization, we applied a balanced 8:8:8 N–P–K chemical fertilizer (Agri-system Co., Ltd), adding approximately 175 g/m^2^ to the Nara field site (fertilized plot), and subsequently quantified soil phosphorus, nitrogen, and potassium concentrations as described in soil analysis. Irrigation frequency depended on weather conditions, typically daily. Mulch is applied to both fields to suppress weed growth. In the Nagano fields, we covered the plants with insect nets to prevent insect damage.

### Soil analysis

Soil nutrient measurement was performed using the EW-THA1J soil analyzer (Air-Water Biodesign Co.) with the EW-T102J cartridge, following the manufacturer’s instructions. Optimal nutrient levels (Fig. S1b) were determined based on the manufacturer’s recommended values.

### Fungal inoculation on agar

Agar plugs of fungal cultures grown on Mathur’s media were transferred to new Mathur’s flasks and incubated at 25°C for 5 days in normal continuous light conditions (10 µmol m^-2^s^-1^). Spores were harvested using DW, washed once, and resuspended in 5-10 mL DW. Spore concentration was determined by loading 10 µL of the suspension into a hemocytometer and counting spores under a light microscope. The suspension was then diluted to 3 μL of 5 × 10^3^ spore/mL before next steps.

Half-strength MS medium (+N) used in this study are based on Gruber et al. (2013) ^46^, with modification (See Table S1). N deficient media was prepared as above with modifications for N0, for N110 and for N275 (See Table S1).

7-day-old *A. thaliana* seedlings were transferred to half-strength MS agar plates with 55 mL nutrient media (120x120x17 mm square plate, Greiner). Fungal spores (3 μL of 5 × 10^3^ spore/mL) were inoculated 3 cm below the seedlings. Plates were placed vertically in a plant growth chamber at 22°C (± 1°C) under fluorescent light (100 µmol m^−2^ s^−1^, 16 h light/8 h dark), around 40% humidity unless otherwise described. To avoid the positional effects, plates were shuffled regularly. The effects on plant growth by fungal inoculation were evaluated by measuring shoot fresh weight or chlorophyll content.

### Leaf chlorophyll measurements

The method of chlorophyll quantification was adapted from the previous report ^47^. 5∼10 mg of crushed leaf tissue from each treatment was sampled and weighed. Chlorophyll was extracted by adding 800 µL chilled cold 100% acetone, and samples were shaken for 10 min until plant tissue was discolored. Centrifuge the tubes at 20,000 x g, 4°C, for 5 minutes and transfer 400 µL of supernatant into another 2 mL tube preloaded with 100 µL of DW and mix using a vortex mixer. After the samples were diluted four times with 80% acetone, the absorbance of tissue-free chlorophyll extract was measured at 646 nm, 663 nm, and 750 nm with an Eppendorf Biospectrometer, following the formula.

- A: Chlorophyll a (µg/ml) = 12.25 × (OD_663_ − OD_750_) − 2.85 × (OD_646_ − OD_750_)
- B: Chlorophyll b (µg/ml) = 20.31 × (OD_646_ − OD_750_) − 4.91 × (OD_663_ − OD_750_)

### RNA extraction and RNA-sequencing analyses

Total RNA was extracted using Nucleospin^®^ RNA Plant (Macherey-Nagel) according to the manufacturer’s instructions. For RNA-seq analysis, ∼ 2 µg of RNA per sample were sent to Rhelixa (Japan) for sequencing. After Poly-A selection by NEBNext Poly(A) mRNA Magnetic Isolation Module, the strand-specific libraries were generated by NEBNext Ultra II Directional RNA Library Prep Kit. The generated libraries were sequenced by Illumina NovaSeq 6000 platform (150 bp x 2 paired-end), resulting in approximately 26.7 M reads per sample for both RNAseq analyses (for Fig. 2-3, Fig. 5) in this study. The obtained reads were mapped to the Ct4 genome or *Arabidopsis* genome (TAIR10) assembly by hisat2 (v2.2.1) ^48^. The obtained sam files were converted to the corresponding bam files by samtools (v1-19) ^49^. featureCounts (v 2.0.6) ^50^ were used to obtain the count data of genes from the bam files. The generated raw counts were used to analyze DEGs by R Bioconductor package DESeq2 after filtering off the genes with less than 10 counts^51^. The following statistical analysis and data visualization were conducted in R packages, such as ggplot2 ^52^. DEGs were identified using an FDR threshold of < 0.05. Transcripts per million (TPM) were then calculated

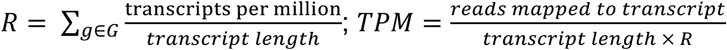

Principal coordinate analysis (PCoA) was plotted based on TPM, using R package ggplot2, and the 95% confidence ellipse was plotted by stat_ellipse function. Heatmaps representing gene expression profiles were generated based on the log_2_ TPM using the R package pheatmap and ComplexHeatmap ^54,55^. GO analysis for *A. thaliana* genes was conducted by g:Profiler, and highlighted terms representing the driver terms (i.e., “leading gene sets that give rise to other significant functions in the neighborhood”) were chosen for further data visualization^56^.

### DNA extraction

Total DNA was extracted from the whole root compartments in soil pot assays using cetyltrimethylammonium bromide (CTAB) method. CTAB extraction buffer was prepared with 3% (w/v) CTAB, 100 mM Tris-HCl (pH 8.0), 1.4 M NaCl, and 20 mM EDTA, dissolved by heating to 60°C. Before use, β-mercaptoethanol was added to a final concentration of 0.2% (v/v). Samples were homogenized using a Shake Master, and 400 µL of pre-warmed CTAB buffer was added to 2 mL tubes containing ground material. Tubes were incubated at 60°C for 30 minutes, with gentle inversion every 10 minutes. After cooling to room temperature, 300 µL chloroform:isoamyl alcohol (24:1) was added, mixed gently, and centrifuged at 12,000 rpm for 3 minutes. The aqueous phase (300 µL) was transferred to 300 µL pre-chilled isopropanol and incubated for 10 minutes. DNA was pelleted by centrifugation at 12,000 rpm, 4°C for 10 minutes, then washed twice with 70% ethanol (centrifuged at 12,000 rpm, 4°C for 5 minutes each). Pellets were air-dried for 30 minutes and resuspended in 50 µL TE buffer (pH 7.5) at 37°C for 15 minutes. DNA concentration was quantified using a NanoDrop One (Thermo) spectrophotometer, and samples were stored at −30°C.

### Quantitative real-time PCR

Total RNA was extracted using Nucleospin^®^ RNA Plant (Macherey-Nagel) according to the manufacturer’s instructions. For the reverse transcription PCR (RT-PCR), cDNA was synthesized from 200 to 500 ng total RNA using the PrimeScript™ RT Master Mix (TaKaRa) in a volume of 10 µL.

cDNA or genomic DNA (gDNA) were then subjected to THUNDERBIRD^®^ Next SYBR qPCR Mix (TOYOBO) with 250 nM primers using CFX Opus 384 Real-Time PCR System (BIORAD) in a volume of 10 µL, or Power™ SYBR™ (Thermo) with 1.6 µM primers using the AriaMx real-time PCR system (Agilent) in a volume of 12 µL. Primers used are listed in Table S9.

### Confocal and light microscopy

Fungal colonization on *A. thaliana* roots was visualized by confocal microscopy (OLYMPUS FLUOVIEW FV3000) using 20.0 3 UPLXAPO20X objective lens (0.8 numerical aperture). Excitation dichroic mirrors DM405/488/561 were used. Variable barrier filters for 500 nm–540 nm were used for the detection of GFP fluorescent signals, and 570 nm–670 nm for mCherry, respectively. The plant cell membrane was labeled by PIP2A (aquaporin) fused with mCherry (PIP2A-mCherry). Ct4 line constitutively expressing GFP (Ct4-GFP) were generated as previously described^29^.

Pam 10 recruitment assay (Fig. 5c, Fig. S10c-d) was conducted using agar system. 7-day-old plants were transferred to agar plates with sufficient N, low N (275 µM), and low N supplemented with amino acids (+N, -N, -N + AA, respectively). Pam10 and Ct4 inoculant concentration was used as co-inoculation assay above. Pam10 colonization on Ct4 hyphae and plant root was observed by Leica S8AP0 stereo microscope (Fig. 5c, Fig. S10d) and Shimadzu BA81-6T-4000XBMH digital microscope using 10x objective lens (Fig. S10c-d).

### Co-culture inhibition assay of Ct and Pam10 in liquid Mathur’s medium

Fungal inhibition by Pam10 in liquid culture was assessed using a modified Mathur’s co-culture assay (Fig. S11h–i). Fungal spores and Pam10 cells were prepared as described above. For each assay, 10 μL of fungal spore suspension (5 × 10^4^ spores/mL) and 10 μL of Pam10 culture (adjusted to OD_600_ = 10^-3^ or 10^-5^) were co-incubated. Fungal biomass was quantified via fluorescence intensity of DsRed (Ct4) or GFP (Ct3) using a Keyence digital microscope (BZ-X1000). Fluorescence was recorded every 10 minutes for 12 hours. The Ct4 lines constitutively expressing DsRed were generated as previously described ^19^, except that GFP was replaced with DsRed in the pBin-GFP-hph vector background.

### Cloning and Agrobacterium-mediated transformation into *A. thaliana* (*AtNRT2.5*)

The primers used for the plasmid construction are listed in Table S9. To construct P35S-NRT2.5, a genomic region of NRT2.5 was PCR amplified using the prime was PCR amplified using the primers NRT2.5-F and NRT2.5-R. The amplified P35S-NRT2.5 sequence was inserted into pENTER^TM^/D-TOPO™ (Invitrogen) using the TOPO reaction to produce an entry clone, known as pENTER^TM^/P35S-NRT2.5. This was then transferred into the destination vector pGWB402, which harbored a 35S promotor sequence, via an LR recombination reaction gateway to produce pGWB402-P35SNRT2.5. For the plant transformation, the constructed plasmid was transformed into *Agrobacterium tumefaciens* (strain GV3101), and an *Arabidopsis* Col-0 was transformed using the floral dip method with the generated Agrobacterium strains^57^. The successful transformations were selected by Kanamycin (50 µg/mL) containing half-strength MS media.

### Generation and validation of *nit2*/*CtΔarea* mutant

The *nit2* mutant was generated using a protocol adapted from Brooker et al. (1991) ^31^ . Agar plugs of Ct4 WT grown on Mathur’s agar were transferred to Mathur’s agar medium amended with potassium chlorate 20 g/L (MC) and incubated for 4-10 days. Hyphal tips from fast-growing sectors, indicative of chlorate-resistant mutants (*nit* mutants), were isolated and transferred to basal medium (BM) supplemented with 2 g/L NaNO_3_. The BM was prepared (per liter) as follows: sucrose, 30 g; KH_2_PO_4_, 1 g; MgSO_4_‧7H_2_O, 0.5 g; agar, 20 g; and trace element solution, 0.2 ml. The trace element solution contained (per 100 ml): citric acid, 5 g; ZnSO_4_‧7H_2_O, 5 g; Fe(NH_4_)_2_(SO_4_)‧6H_2_O, 1 g; CuSO_4_‧5H_2_O, 0.25 g; MnSO_4_‧H_2_O, 50 mg; H_3_BO_4_, 50 mg; and NaMoO_4_‧2H_2_O, 50 mg. After 5 days of incubation, isolates exhibited reduced growth in BM+NO_3_, as exemplified in Fig. 3c, were selected as *nit* mutant candidates. These candidates were transferred to screening media consisting of BM supplemented with one of the following N sources: 0.5 g/L sodium nitrite, 1 g/L ammonium tartrate, 0.2 g/L uric acid, or 0.2 g/L hypoxanthine. A *nit2* mutant candidate was identified based on its impaired growth on all media except partial recovery on ammonium, indicative of a defect in global N utilization regulation.

The *nit2* candidate was validated by sequencing the *CtAREA* genomic region of *Colletotrichum tofieldiae* MAFF 712334 (Ct4: GenBank ID: GCA_022836575.1). Ct4 wild-type and *nit2*/*CtΔarea* candidate strains were cultured for spore production as described above. Fungal spores (10 μL of 5 x 10^3^ spores/mL) were inoculated into 20 mL liquid Mathur’s medium for 2-3 days, mycelia were then harvested and homogenized using a Shake Master.

Genomic DNA was extracted using the Chelex: fungal mycelia were resuspended in 100 µL of 10% (v/v) Chelex^®^ -100 (BIORAD) in 10 mM Tris-HCl (pH 8.0) and vortexed. Tubes were subjected to three freeze– boil cycles (1 min in liquid N_2_ followed by 1 min at 100°C), then incubated at 100°C for 5 min and vortexed for 30 s. Lysates were centrifuged at 12,000 rpm for 6 min at 4°C, and 20-40 µL of the supernatant was collected. DNA concentration was quantified using a NanoDrop One (Thermo) spectrophotometer, and gDNA was stored at –30°C or used directly for PCR.

The full-length *CtAREA* locus (∼3,700 bp) was amplified using KOD FX Neo PCR Master Mix (TOYOBO), and PCR products were validated by agarose gel electrophoresis. The target band was extracted using the FastGene™ Gel/PCR Extraction Kit and further used for amplification of approximately eight overlapping fragments. These fragments were subjected to Sanger sequencing, and resulting sequences were aligned and compared between Ct4 WT and mutant candidates to confirm the presence of mutations in the *CtAREA* region. Primers used are listed in Table S9.

### Fungal growth assay in Mathur’s media (vitro growth)

Equal-sized agar plugs of ten-day-old fungal cultures grown on Mathur’s media were transferred to new BM plates and incubated at 25°C. Fungal growth was determined five days after incubation by visualizing and measuring the colony radius from center to edge. Fungal spore counts were conducted as described above.

### 15N translocating assays

A two-compartment agar plate system was established (Fig. 4a). Both compartments contained half-strength MS medium, with modification applied exclusively to the hyphal compartment (HC), which was inaccessible to plant roots. In the HC, ^15^N-labeled potassium nitrate (K^15^NO_3_) was used to replace the standard K^14^NO_3_. This HC compartment will be inoculated with Ct4 or *CtΔarea* mutants spore suspension or controls, which is either DW (mock) or the pathogenic fungus *Colletotrichum incanum* (Ci). The plant-hyphal compartment (PHC) was supplemented with N at three levels: none (N0), low (N110; 110 µM), or sufficient (11,400 µM) (See Table S1). After the inoculated fungi have fully spread to the PC from the HC, the HC will be removed to avoid any possible ^15^N contamination. Subsequently, 7-day-old seedlings will be transferred to the PHC. Plates were placed vertically in a plant growth chamber at 22°C (± 1 °C) under fluorescent light (100 µmol m^−2^ s^−1^, 16 h light/8 h dark), around 40% humidity.

At 20 dpi, shoots were harvested, dried at 60°C for 48 h, and ground to a fine powder. Grounded sample was packed into Ø 5×9 mm tin capsules (24006400, Thermo) and analyzed for ^15^N atom% using an isotope ratio mass spectrometer (DELTA V Advantage, Thermo). Alanine was used as a working standard for correction. As ^15^N is a natural isotope, even in the PHC compartment where ^15^NO_3_ was not supplied, ^15^N made up around 0.365 atom%. Hence, to trace the ^15^N translocation, we traced the increased in ^15^N atom% in Ct4-treated plants.

### Bacterial genome sequencing and comparative genomic analyses

Bacterial DNA was extracted using the NucleoSpin^®^ Microbial DNA Kit (Macherey-Nagel (TaKaRa)) according to the manufacturer’s standard protocol. Subsequent procedures, up to and including gene prediction, were carried out by BGI with standard procedures. Briefly, the extracted DNA was used for library preparation for PacBio sequencing. For genome assembly, subreads were error-corrected using proofread (v2.12) and assembled with Falcon (v0.3.0) ^58^ under the following parameters: -v -dal8 -t32 -h60 -e.96 -l500 -s100 -H3000. Single-base errors in the assembled sequences were further corrected using DNBseq data. Gene prediction was performed with Prokka ^59^ and RAST ^60^ using the default parameters then manually curated based on NCBI’s Conserved Domain Database (CDD) ^61^.

The KEGG annotation was performed using KofamScan (v1.3.0) with the February 2025 release of the KOfam HMM profiles ^62^. KofamScan used hmmsearch from the HMMER suite (v3.4) to align proteins to the KOfam profiles ^63^. Each protein sequence with a significant KofamScan hit was associated to the metabolic pathway assigned by KofamScan, and multiple heatmaps were generated comparing different metabolic pathways for the bacterial strains under investigation. The Python code to perform the gene prediction, KEGG annotation, and to generate the heatmaps is available at https://gitlab.com/salvo981/genome_data_analysis.

### Meta-analysis of bacterial 16S rRNA gene amplicon

The generated paired-end reads were trimmed, quality-filtered, and denoised, and the sequences with ≥99% similarity were sorted into amplicon sequence variants (ASVs) using the DADA2 v1.18 program package ^64^. As described previously by Utami et al. (2019) ^65^, the procedure was carried out with modifications to the trim-left parameters set to 19 and 18 to remove the forward and reverse primer regions. The obtained ASVs were phylogenetically classified using SINA v.1.2.11 ^66^ with the SILVA v138 database, and those not assigned to Bacteria were discarded. Statistical analyses of soil microbial diversity were carried out using QIIME^67^. Beta diversity analysis was performed based on the Bray-Curtis distance matrix and plotted as PCoA by the R package ggplot2 v3.5.2. Differences in beta diversity were tested using permutational analysis of variance (PERMANOVA; 999 permutations).

For deciding specifically enriched ASVs, the generated raw reads were used to analyze differentially enriched by R Bioconductor package ALDEx2 (v1.34.0) after filtering off the ASVs with 0 counts in all treatments, using mock-treated samples as preference. Only ASVs that were enriched under N deficiency but not sufficiency were chosen for heatmap. The mean relative abundance and subsequent z-score of the mean relative abundance of these ASVs were calculated using the “scale()” function of R Bioconductor. The heatmap was created using the R package pheatmap and ComplexHeatmap.

### Bacterial inoculation on agar and soil

Bacterial strains were cultured in Luria-Bertani (LB) media composed of tryptone (10 g/L), NaCl (10 g/L), and yeast extract (5 g/L) (final pH = 7.0). 1% (w/v) agar (BD Difco™ Agar) was added for LB agar media.

Bacterial strains were initially cultured on LB agar plates at room temperature for 2–3 days, then sub-cultured into 4 mL LB liquid broth (Greiner Bio-One 14 mL tube) at 25°C for 24 h. Cells were harvested by centrifugation at 5000 × g for 1 min, resuspended in 4 mL DW, and adjusted to OD_600_ = 1. DW was used as the blank. The bacterial suspension was further diluted 10⁴-fold for final inoculants.

For axenic agar assays, N deficient media (N275) was prepared as above with amino acids supplementation as follows: 1.7 nM L-Valine, 1.5 nM L-Leucine, 2.4 nM L-Phenylalanine. Bacterial inoculation was performed by applying either 3 µL droplets or streaking 10 µL of the bacterial inoculant approximately 2 cm below seedlings. For co-inoculation experiments, fungal inoculant Ct4 was added 3 cm below seedlings as previously described. Growth conditions followed those detailed above.

For soil-pot assays, 10 mL bacterial inoculant was directly applied to pots immediately after seedling transfer. Subsequent growth conditions, watering irrigation, and nutrient supplementation followed the fungal inoculation protocol for pot and field assays.

### Evaluation of Pam10 growth by measuring colony size

The N-deficient medium supplemented with amino acids (−N + AA) was prepared as described above, with a slight modification in amino acid concentrations (AA: 1.7 nM L-valine, 1.5 nM L-leucine, 1.2 nM L-phenylalanine. Pam10 cells were harvested as described above and adjusted to OD_600_ = 0.1. The bacterial suspension was further diluted 10⁴-fold for final inoculants. 10 µL of bacterial suspension was spread on agar plates with 20 mL nutrient media (Ø 90×15 mm round plate, LABTAS+). Images of the cultured plates (Fig. S10f) were acquired using a SONY α7Ⅳ digital camera (SONY Group Corporation, Japan), and the colony areas of Pam10 were measured using an ImageJ in-house macro. To exclude the edge of Petri dish and the ruler, only the dish area including the region of interest was manually selected and cropped. The images were converted from RGB to 8-bit grayscale, and background subtraction was performed using a rolling ball algorithm with a radius of 50 pixels. Otsu’s method was applied to automatically set the threshold, and binary masks were generated from the thresholder images. Tentative colony areas were calculated for the particles with and area ≧ 200 pixels and circularity between 0.3 to 1.00. Finally, mis-extracted colonies were manually checked and excluded.

## Supplementary Figures

**Figure S1:**
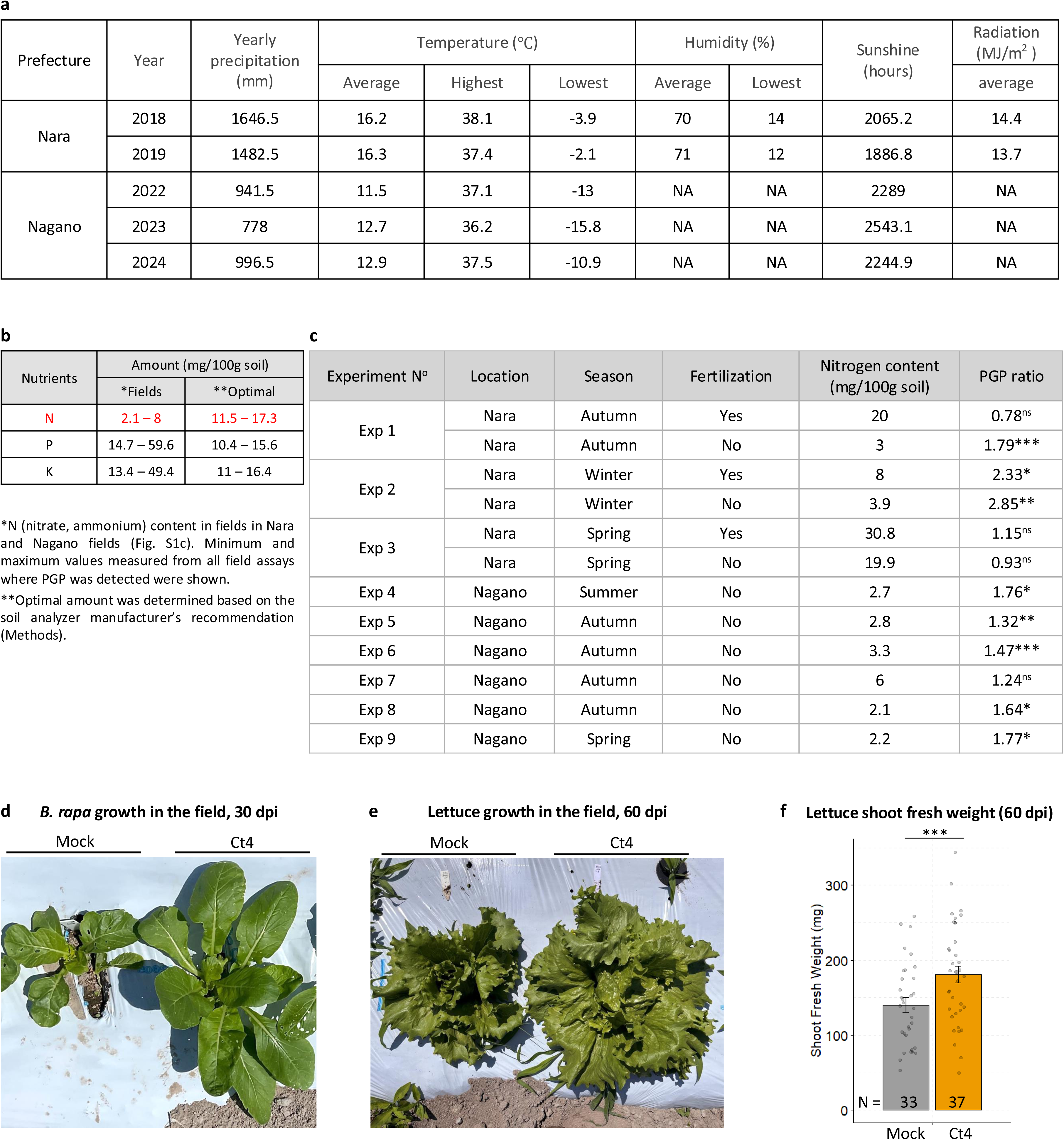
Ct4 enhances *B. rapa* (Komatsuna) and *L. sativa* (lettuce) growth in the N-limiting fields with varied climatic conditions. (a) Nara and Nagano climatic conditions in years that field assays were conducted. Data was retrieved from https://www.data.jma.go.jp/. (b) Nitrogen, phosphorus, potassium nutrient contents in field assays in Nara and Nagano Prefecture. (c) Fertilization status and measured N level in field assays. PGP ratio = Average SFW_Ct4-treated plants_ / Average SFW_Mock-treated plants_. n.s., not significant; *p < 0.05; **p < 0.01; *** p < 0.001, two-tailed t-test. (d) Ct4 enhances *B. rapa* growth in fields. Photos taken at harvest day (Exp 9, 30 dpi). (e-f) Ct4 enhances *L. satica* (lettuce) growth in Nagano fields. Photos taken at harvest day (60 dpi). *** p < 0.001, two-tailed t-test.

**Figure S2:**
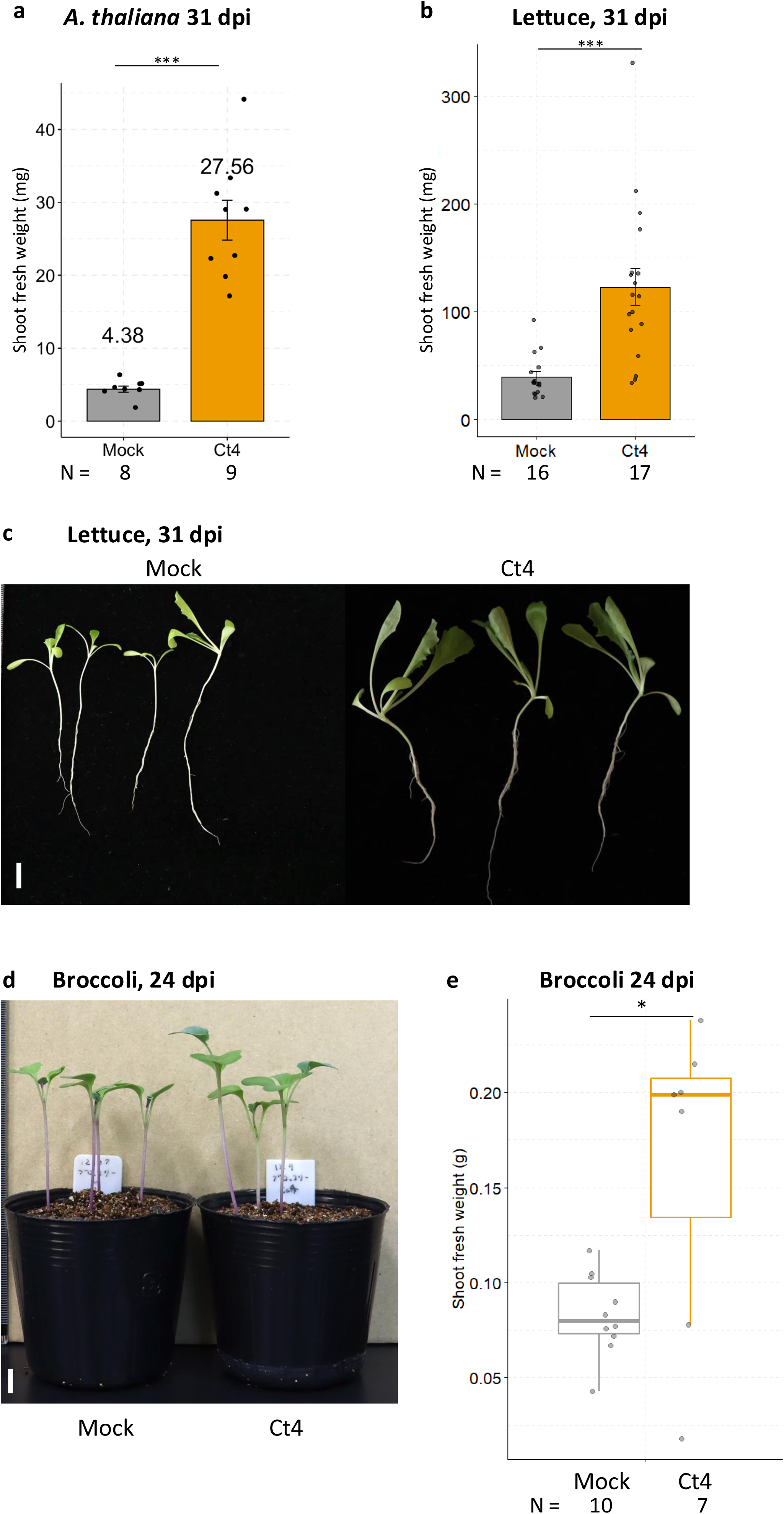
Ct4 enhances the growth of *A. thaliana*, lettuce, and broccoli under N-limiting conditions (110 µM of N supplied) in soil pot assays. (a) Ct4 enhances *A. thaliana* growth in soil pot at 31 dpi. (b-c) Ct4 enhances *L. sativa* (lettuce) growth in soil pot at 31 dpi. Bars = 1 cm. (d-e) Ct4 enhances broccoli growth in soil pot at 24 dpi. Bars = 1 cm. *p < 0.05, *** p < 0.001, two-tailed t-test.

**Figure S3:**
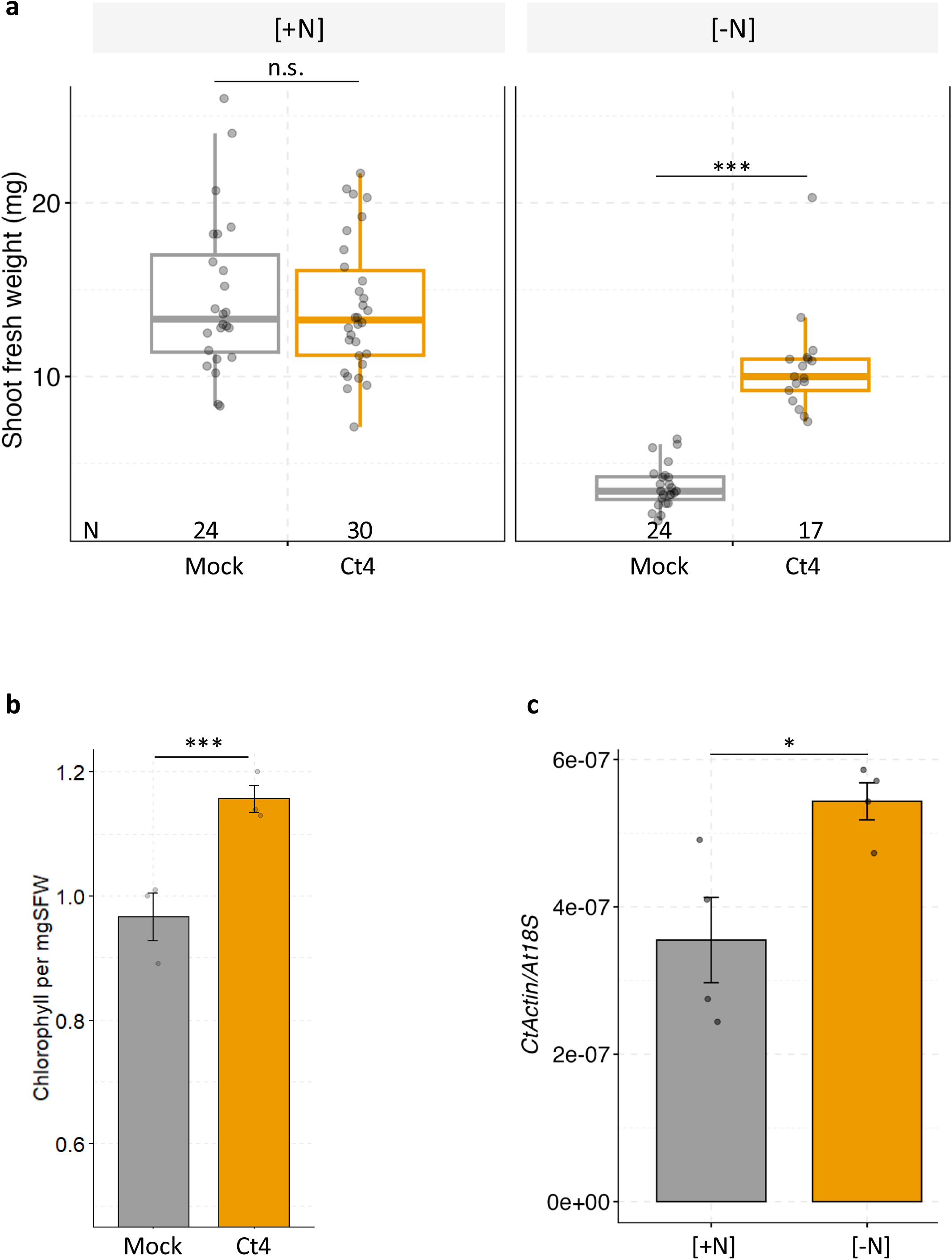
Ct4 enhances *A. thaliana* growth specifically under N-limiting conditions. (a) Shoot fresh weight of *A. thaliana* at 24 dpi in a gnotobiotic agar-plate assay under [+N] and [-N] (N110) conditions. (b) Chlorophyll content in shoots of *A. thaliana* inoculated with distilled water (mock) or Ct4. N = 3. (c) Fungal biomass in roots under +N and –N conditions in (a). N = 4. n.s., not significant; *p < 0.05 ; ***p < 0.001; two-tailed t-test.

**Figure S4:**
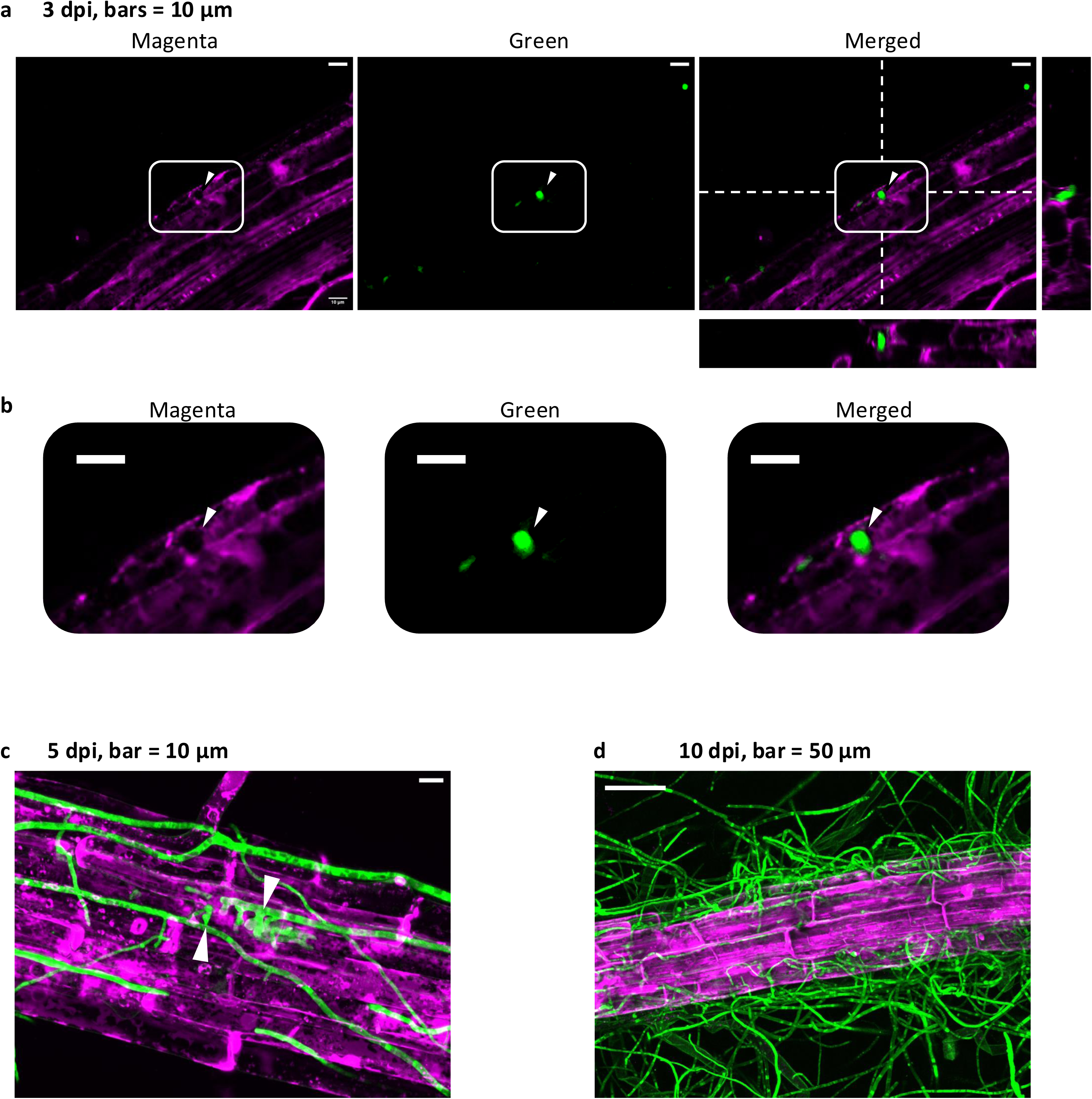
Confocal microscopic images of Ct4 expressing cytoplasmic GFP (green) and root cells of *A. thaliana* membrane-expressing PIP2A-mCherry (magenta) under N-deficient conditions (N110). Maximum projection of z-stack images were shown. (a-b) Ct4 hypha penetrating a root epidermal cell (arrowheads) at 3 dpi. Representatives of hyphal penetrations are shown as orthogonal views made from areas indicated by white dotted line in (a) and enlarged projected images (b). Bar = 10 μm. (c) Ct4 hyphae elongated and branched within colonized plant cells (arrowheads) at 5 dpi. (d) Extensive Ct4 colonization was observed on the root at 10 dpi.

**Figure S5:**
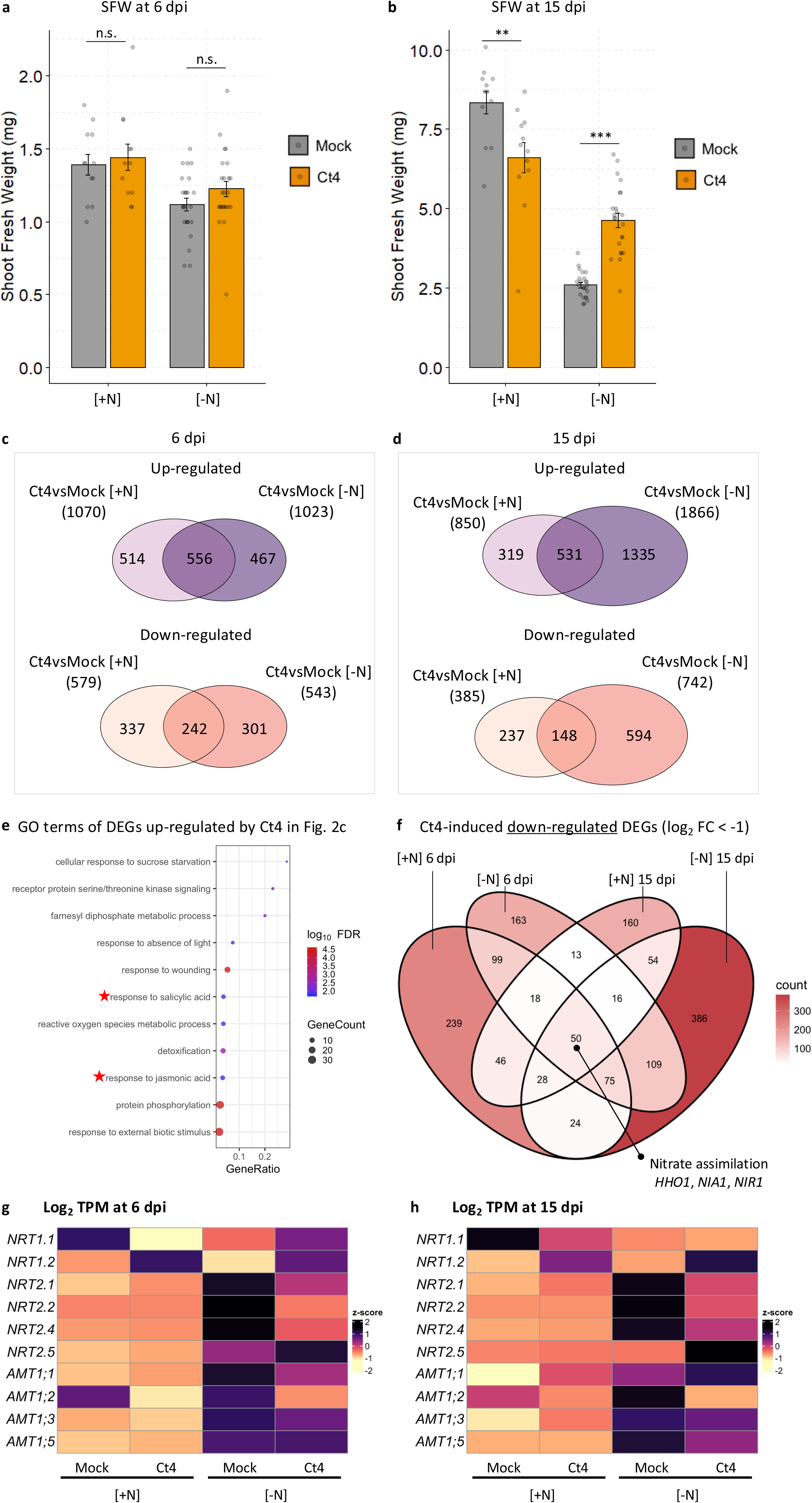
RNA-Sequencing analysis during *A. thaliana* root colonization by Ct4 under N-sufficient [+N] and N-deficient [-N] (N110) conditions. (a-b) SFW (mg) of *A. thaliana* at 6 dpi (a) and 15 dpi (b). n.s., not significant; **p < 0.01 ; ***p < 0.001; two-tailed t-test. (c-d) DEGs up-regulated (log_2_ FC > 1, FDR < 0.05) or down-regulated (log_2_ FC < -1, FDR < 0.05) by Ct4 at 6 dpi (c) or 15 dpi (d) under [+N] or [-N]. (e) GO terms of 242 DEGs up-regulated by Ct4 in all conditions from Fig. 2c. (f) Ct4-induced down-regulated DEGs (log_2_ FC < -1, FDR < 0.05) at 6 dpi and 15 dpi under [+N] or [-N]. (g-h) log_2_ TPM of *AtNRT*s and *AtAMT*s at 6 dpi (g) or 15 dpi (h).

**Figure S6:**
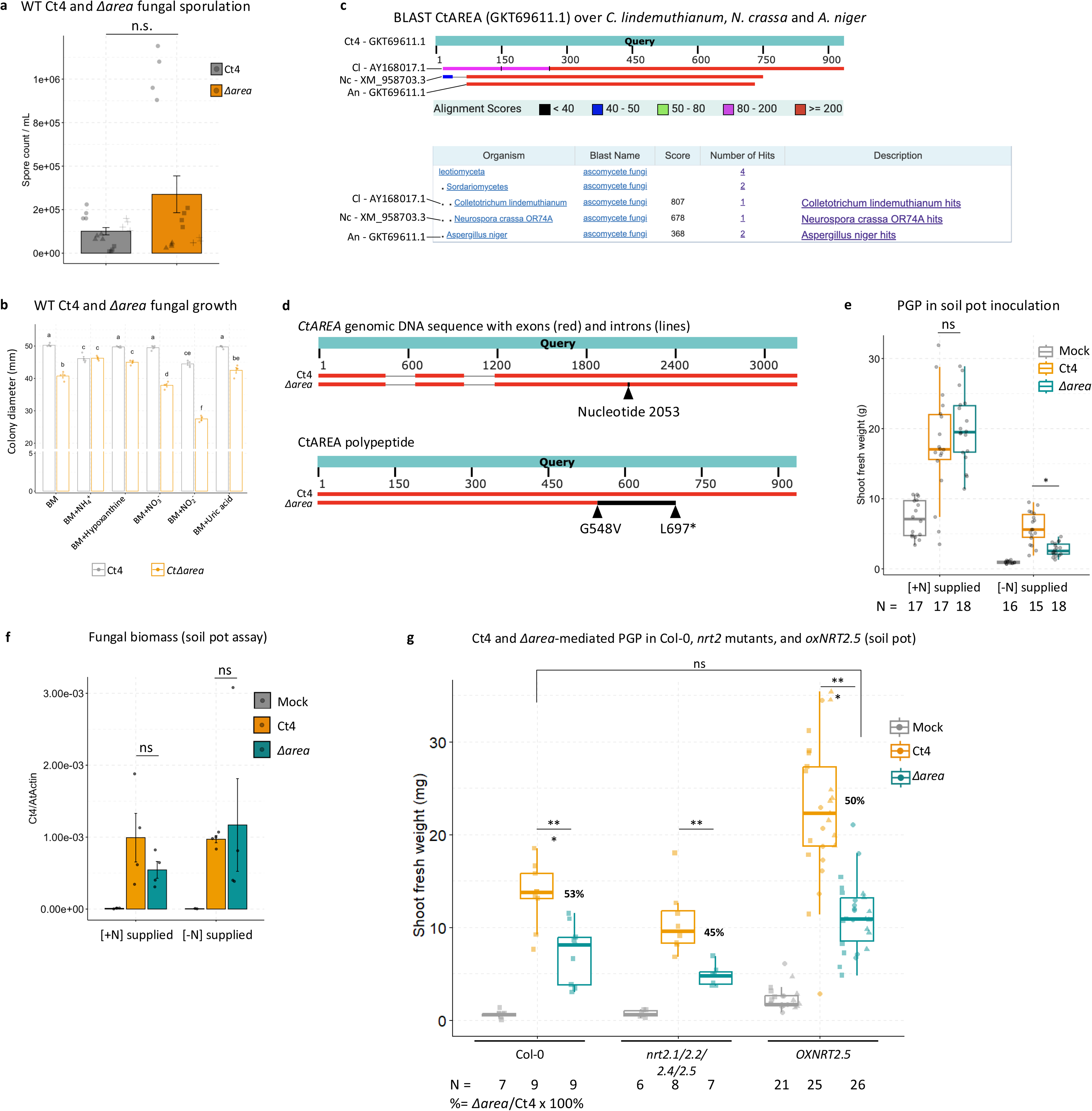
*nit2*/*CtΔarea* mutant defective in nitrate utilization and PGP. (a) Spore count of Ct4 WT and *nit2*/*CtΔarea* mutant grown in Mathur’s medium. n.s., not significant, two-tailed t-test. (b) Colony diameter Ct4 WT and *nit2*/*CtΔarea* mutant in basal media (BM) and BM supplemented with indicated N sources. Letters: post-hoc Tukey, p < 0.05. (c) Protein BLAST (NCBI) of Ct4AREA (GenBank: GKT69611.1) against AREA/NIT2 orthologs in *C. lindemuthianum*, *N. crassa* and *A. niger*. (d) Sanger sequencing of the *CtAREA* genomic locus showing the mutation site in *CtΔarea* (upper panel, arrow), leading to a frameshift from G548V and premature stop at L697* (lower panel, black region). (e) Shoot fresh weight (mg) of *A. thaliana* (soil pot assay) treated with mock, Ct4, or *CtΔarea*. Soil pots were irrigated with ½-strength MS solution containing 11.4 mM ([+N] supplied) or 110 µM ([-N] supplied) of total N. n.s., not significant; *p < 0.05, two-tailed t-test. (f) Ct4 biomass in roots from (e), quantified by qPCR using Ct4-specific primers (Table S9) and normalized to *A. thaliana* Actin. n.s., not significant, two-tailed t-test. (e) Shoot fresh weight (mg) of *A. thaliana* (soil pot assay) treated with mock, Ct4, or *CtΔarea*. Soil pots were irrigated with ½-strength MS solution containing 110 µM of total N. Different OX lines were shown by square, round, and triangle symbols. n.s., not significant; **p < 0.01, two-tailed t-test.

**Figure S7:**
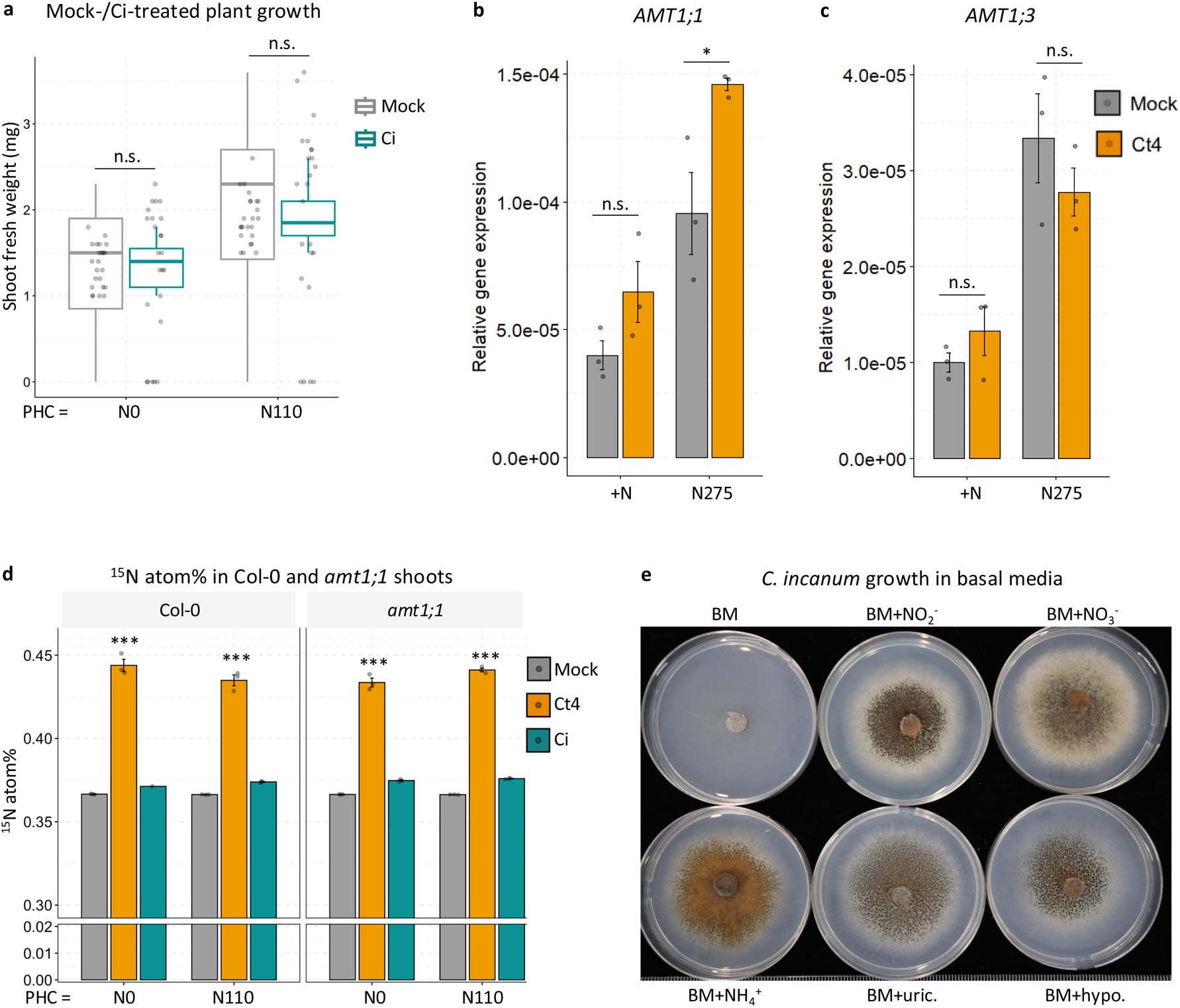
Ct4, but not Ci, transfers N independently of plant NRT2s and AMT1;1. (a) Shoot fresh weight (mg) of *A. thaliana* in N-limiting (agar) conditions treated with mock or Ci. (b-c) Relative gene expression of *AtAMT1;1* and *AtAMT1;3* in Ct4-inoculated roots under N-sufficiency and N-deficiency, normalized to At18S. N = 3. (d) ^15^N atom% ratio in shoots of *A. thaliana* Col-0 or *amt1;1* plants treated with Mock, Ct4, or Ci. HC was supplemented with 110 µM of N, PHC was not supplemented with N (N0) or supplemented with 110 µM of N (N110). N = 3. (e) Ci growth on basal media BM or BM supplemented with nitrite, nitrate, ammonium, uric acid (BM+uric.), and hypoxanthine (BM+hypo.). Representative plates were shown. N = 3. n.s., not significant, *p < 0.05, ***p < 0.01, two-tailed t-test.

**Figure S8:**
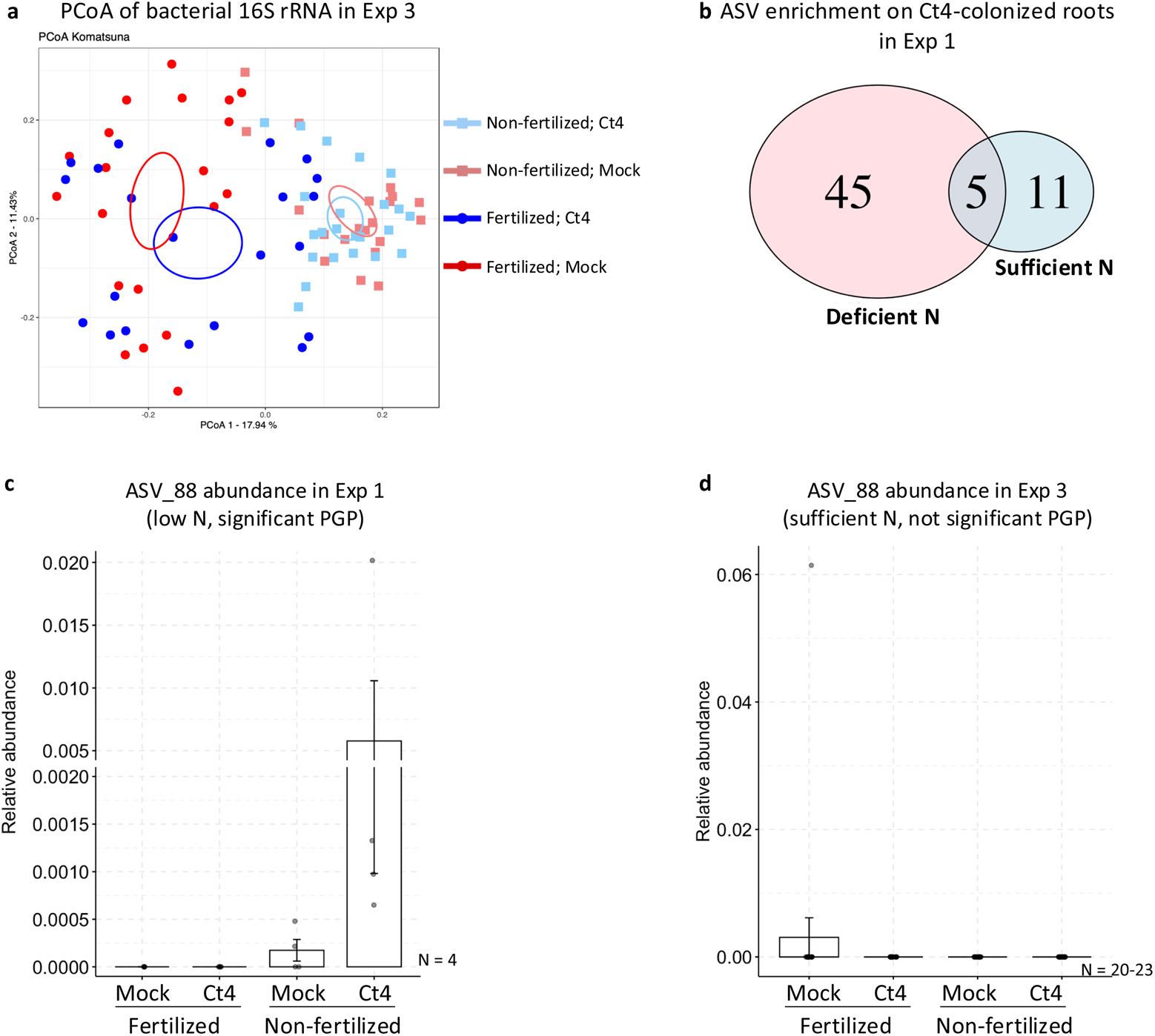
Meta-analysis of bacterial 16S rRNA gene amplicon on Ct4-colonized *B. rapa* roots in field assays (Nara). (a) PCoA of *B. rapa* root bacterial community profiles in fertilized and non-fertilized fields in Exp 3 (Fig. S1c), where N was sufficient in both fields. p < 0.05, ANOSIM. (b) Number of bacterial amplicon sequence variants (ASVs) significantly enriched in Ct4-inoculated roots under fertilized (Sufficient N) and non-fertilized (Deficient N) fields in Exp 1 (Fig. S1c), as identified by ALDEx2 (adjusted p < 0.05). (c-d) Relative abundance of ASV_88 (Pam10) in Ct4-inoculated roots under field conditions in Fig. S1c: (c) N-deficient, non-fertilized soil (Exp 1) and (d) N-sufficient conditions in fertilized and non-fertilized soils (Exp 3).

**Figure S9:**
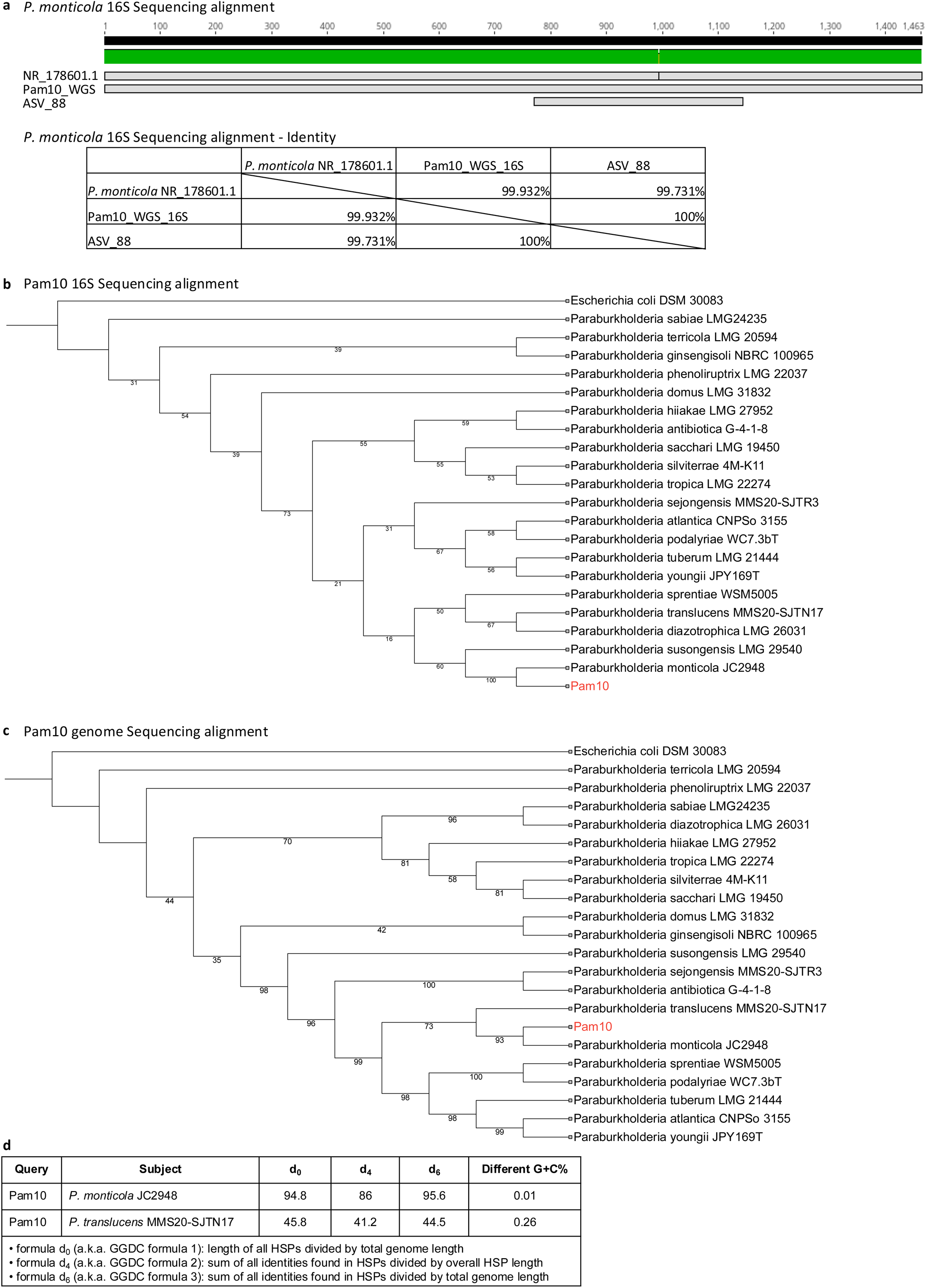
Pam10 (ASV_88) taxonomic identification at the species level. (a) Alignment of 16S rRNA gene sequences of Pam10. Shown are the NCBI sequence of *Pseudomonas monticola* (NR_178601.1), the 16S region obtained from Pam10 whole-genome sequencing (Pam10_WGS_16S), and the corresponding ASV from 16S amplicon analysis (ASV_88). (b) 16S rRNA-based taxonomy for Pam10 using TYGS (https://tygs.dsmz.de) showing high 16S rRNA gene similarity between the in-house isolated Pam10 and *P. monticola*. (c) Genome-based taxonomy for Pam10 by Genome-to-Genome Distance Calculator method via TYGS (https://tygs.dsmz.de) showing high genomic similarity between the in-house isolated Pam10 and *P. monticola*. (d) Represent result from TYGS for genome-based taxonomy in (c) for highest and second highest strains to Pam10 (*P. monticola* and *P. translucens*, respectively).

**Figure S10:**
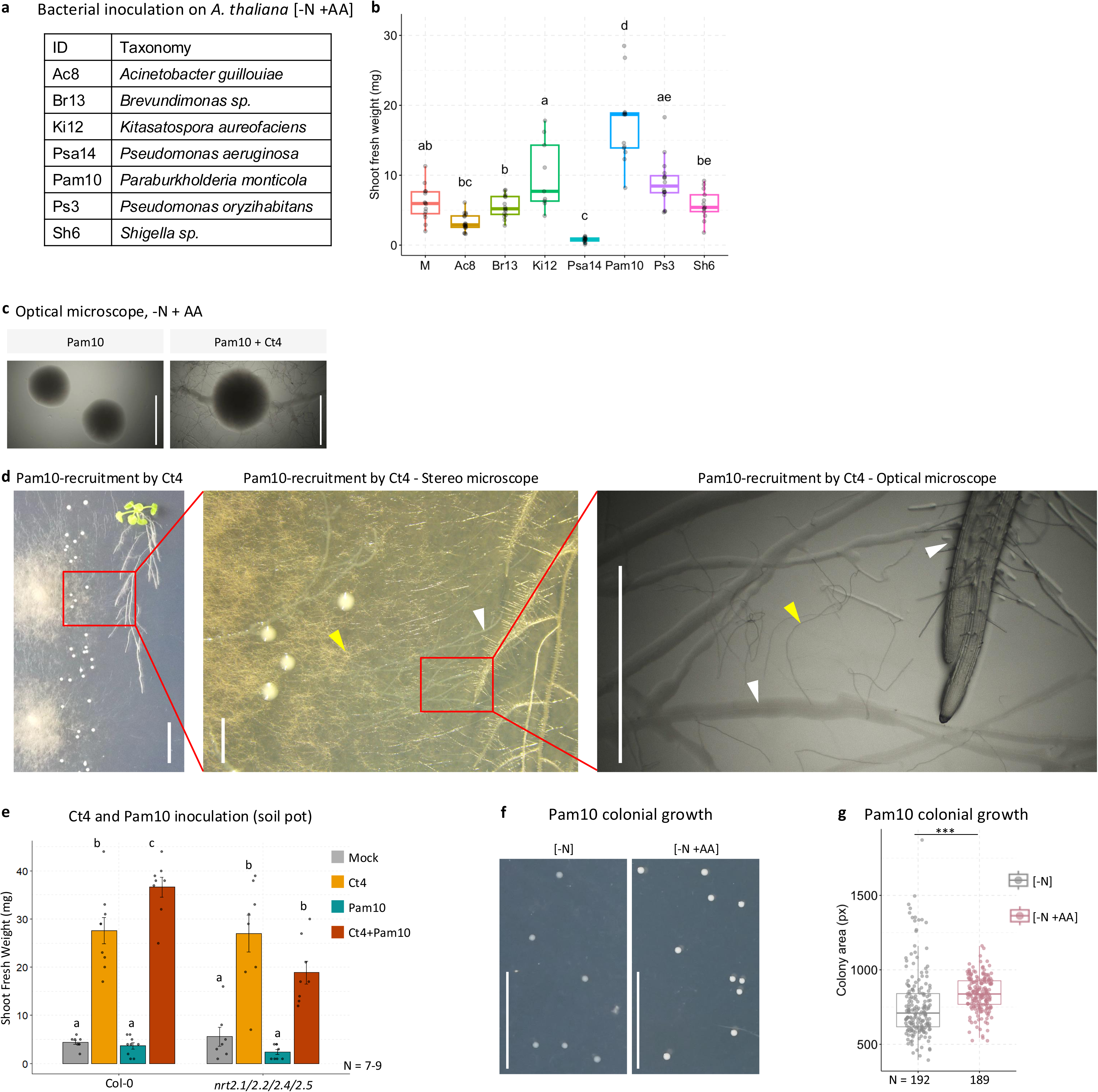
Pam10 rides fungal hyphae and approach plant root, subsequently promote plant growth when amino acids were supplemented. (a-b) Bacterial strains isolated from *A. thaliana* roots in soil pot assays (left), and their effects on *A. thaliana* shoot growth at 26 dpi under N-deficient conditions supplemented with amino acids (right). Letters: post-hoc Tukey, p < 0.05. (c) Optical microscopy images of Pam10 colonies grown alone (left) or in co-culture with Ct4 (right) under N-limited conditions supplemented with amino acids (–N + AA). (d) Ct4-induced recruitment of Pam10 to plant roots under –N + AA conditions. Images shown from left to right with scale bars: 1 cm, 0.2 cm, and 0.1 cm. White arrows: Pam10 biofilm; yellow arrow: Ct4 hyphae. (e) Single and co-inoculation assays of Ct4 and Pam10 in *A. thalian*a Col-0 and *nrt2.1/2.2/2.4/2.5* mutants under N-limiting conditions in soil pots. Letters: post-hoc Tukey, p < 0.05. (f) Pam10 growth in N-deficiency with/without amino acids supplementation ([-N] and [-N +AA], respectively). Bar = 1 cm. (g) Colony size of Pam10 grown in (f). ***p < 0.01, two-tailed t-test.

**Figure S11:**
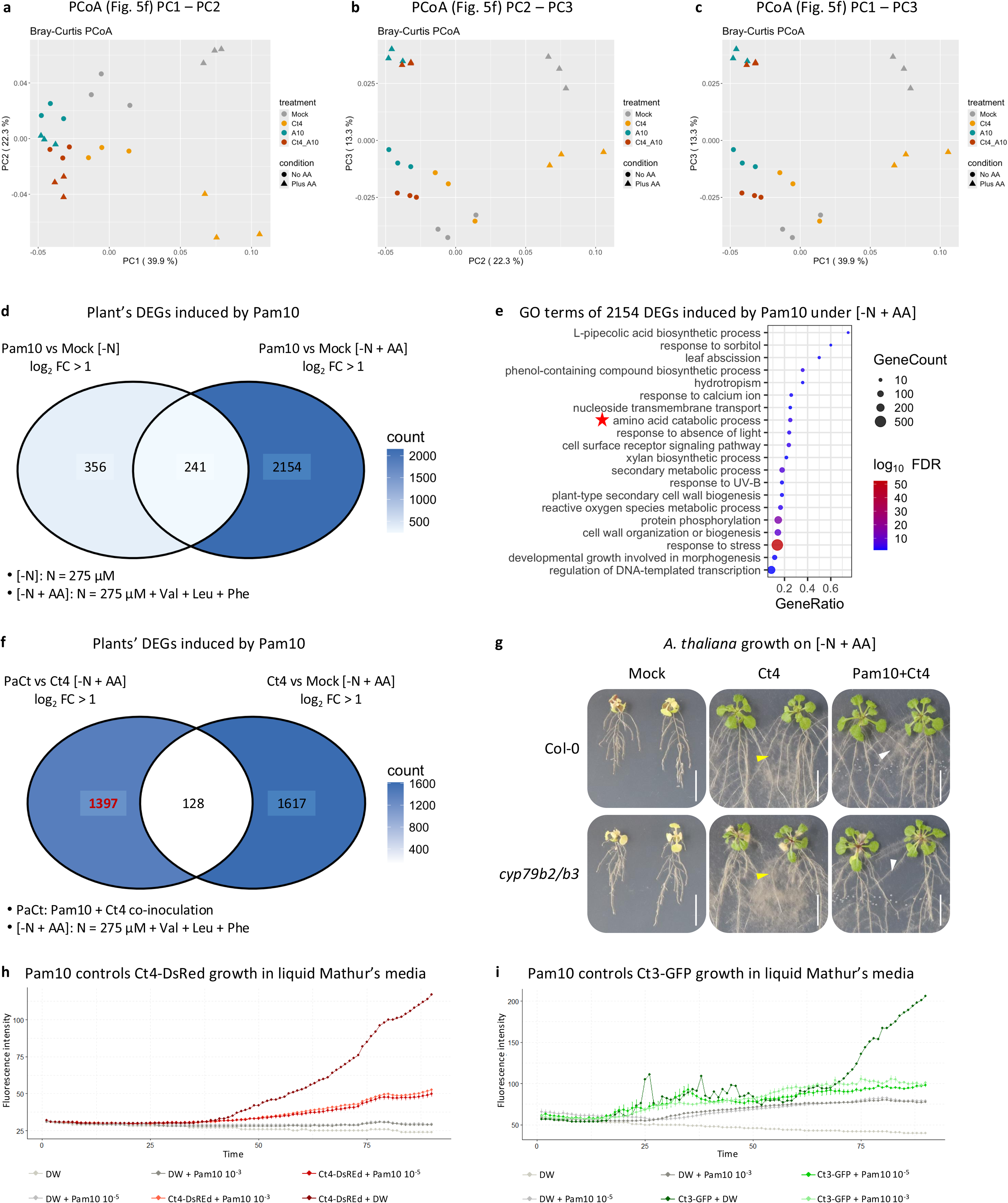
Transcriptomic responses and interactions between Ct4, Pam10, and *A. thaliana* under N-limited conditions (N275) supplemented with amino acids. (a–c) Principal coordinate analysis (PCoA) plots based on transcript per million (TPM) expression values for *A. thaliana* plants shown in Fig. 5f. (d) Venn diagram showing the number of DEGs induced by Pam10 under N deficiency (–N) and N deficiency supplemented with amino acids (–N + AA) at 17 dpi (log₂ FC > 1, FDR < 0.05). (e) Gene Ontology (GO) term enrichment analysis of 2154 *A. thaliana* genes specifically upregulated by Pam10 under [–N + AA] conditions identified in (d). (f) Venn diagram showing the number of *A. thaliana* DEGs in response to Ct4 + Pam10 co-inoculation vs. Ct4 single inoculation, and Ct4 vs. mock under –N + AA conditions at 17 dpi (log₂ FC > 1, FDR < 0.05). 1397 DEGs were specific to the PaCt co-inoculation and were showed in Fig. 5g heatmap. (g) Single and co-inoculation assays of Ct4 and Pam10 in *A. thaliana* Col-0 and *cyp79b2/b3* double mutants under N-limiting conditions supplemented with amino acids in agar system. White arrows: Pam10 colonies; yellow arrow: Ct4 hyphae. (h-i) Pam10 controls fungal growth of Ct4 and Ct3 in liquid Mathur’s medium, as quantified by fluorescence intensity. Fungal biomass of Ct4-DsRed (h) and Ct3-GFP (i) was measured in co-culture with Pam10 at two bacterial inoculum concentrations: OD_600_ = 10^-3^ and 10^-5^ (10 μL). Fungal spores were inoculated at 5 × 10^4^ spores/mL (10 μL). Fluorescence signals were recorded every 10 minutes for 12 hours. Y-scale: fluorescence intensity, X-scale: time point T, ΔT = 10 minutes.

